# Coronary Artery Disease Transcriptomics Reveals Two Drivers of the Endothelial Cell SR-BI Expression and LDL Transport that Underlie Atherosclerosis

**DOI:** 10.64898/2026.08.05.743006

**Authors:** Linzhang Huang, Yibin Huang, Jiayu Zhu, Jun Peng, Kenian Chen, Ken Chambliss, Qinbo Zhou, Ryan Vela, Dennis Burns, Bo Li, Matthias Peltz, Yun Fang, Lin Xu, Chieko Mineo, Philip W. Shaul

## Abstract

Atherosclerosis is initiated by circulating low-density lipoprotein (LDL) cholesterol transfer into the artery wall, which is mediated by scavenger receptor class B, type I (SR-BI) in endothelial cells(1). Employing single-cell RNA sequencing in human coronary artery disease (CAD) samples, here we show that endothelial SR-BI expression is increased in atheroma, and in endothelial cells with a transcript signature indicative of responding to disturbed blood flow. In vivo in mice hypercholesterolemia and disturbed blood flow independently upregulate endothelial SR-BI; the flow-related upregulation initiates endothelial cell LDL uptake and atherogenesis. Guided by transcription factor networks, it is revealed that HIF-1α binding to human Scarb1 Intron 1 governs endothelial SR-BI transcription, and in mice HIF-1α drives hypercholesterolemia-related SR-BI upregulation and artery LDL uptake. Thus, the two major instigators of atherosclerotic lesion formation, hypercholesterolemia and disturbed blood flow, both upregulate endothelial SR-BI to drive the LDL transport that underlies the disorder. Targeting the processes regulating endothelial SR-BI potentially represents a new therapeutic strategy against CAD.

---

Atherogenesis entails the transport of circulating LDL cholesterol to the subendothelial space, the recruitment of immune cells including monocytes to the artery wall which become macrophages in the intima, and LDL engulfment by the macrophages and vascular smooth muscle cells that take on phagocytic capabilities(2, 3). Single-cell RNA sequencing (scRNA-seq) and assay for transposase-accessible chromatin using sequencing (ATAC-seq) studies in atherosclerotic carotid and coronary arteries have revealed the epigenetic landscape and gene expression present in vascular cells in atherosclerosis in humans(4, 5). However, the dynamics of gene expression and the underlying regulatory processes operative during atherogenesis in humans are poorly understood. Here we bridge these critical knowledge gaps by performing scRNA-seq in coronary artery segments containing versus lacking an overt atheroma from five cardiac transplant subjects with atherosclerosis (Fig. 1a, Supplementary Table 1). The specific impetus was to probe the biology of scavenger receptor class B, type I (SR-BI, encoded by Scarb1), which mediates the endothelial cell LDL transcytosis that drives atherogenesis(1), in human coronary artery endothelium. The data obtained also provide insights into other important aspects of endothelial cell biology and non-endothelial cell gene regulation in the critical context of human coronary artery disease (CAD).

**Fig. 1:**
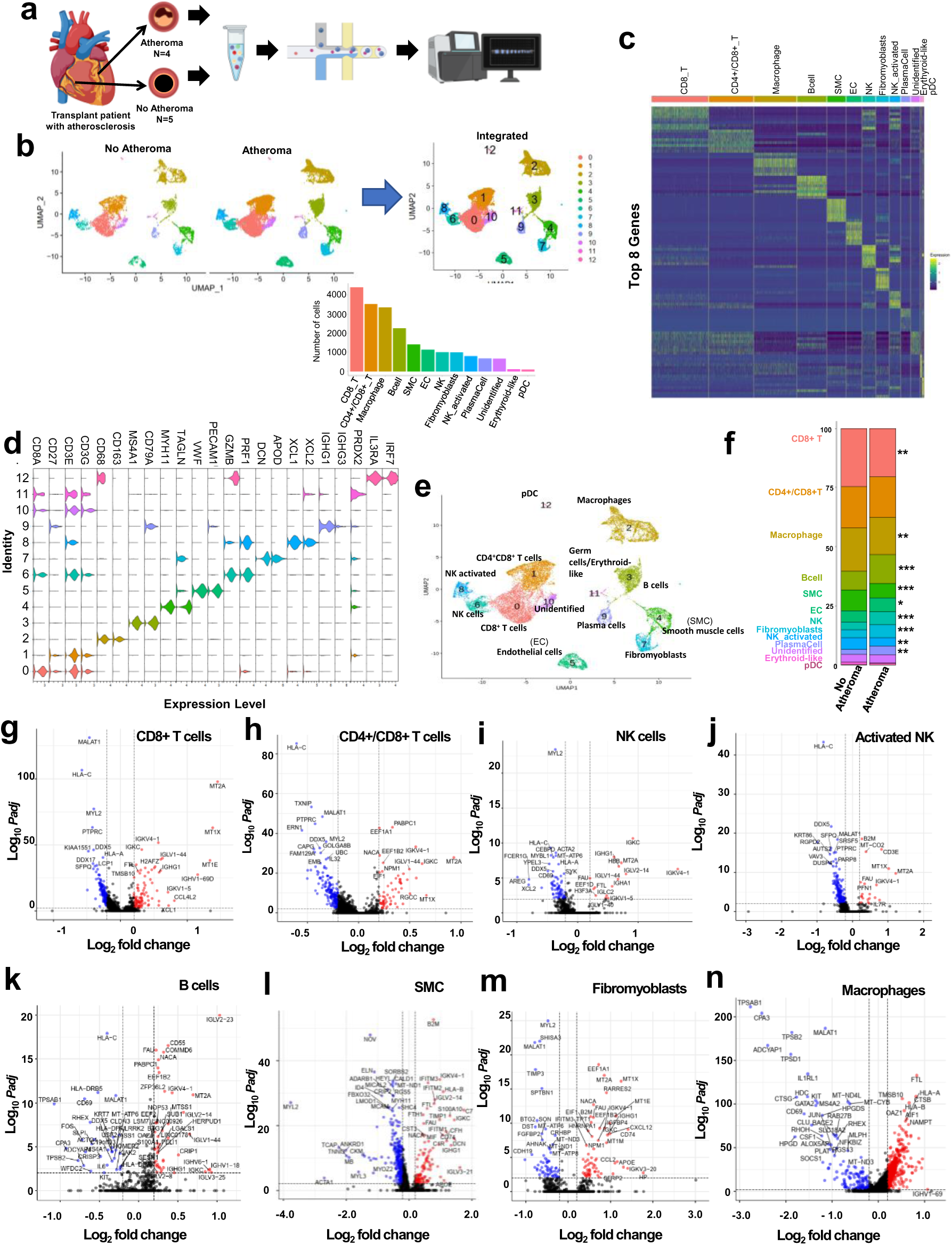
Vascular cell populations and their transcriptomes change with atheroma formation. a, Following the removal of adventitia, scRNA-seq was performed on coronary artery segments containing versus lacking an overt atheroma from five transplant subjects with atherosclerosis. **b,** UMAP plots depicting the 13 cell clusters that were identified, and the number of cells. **c**, Heat map displaying top 8 most abundant genes in each cell cluster. **d**, Marker genes for the cell clusters. **e,f**. UMAP plot indicating cell type identities (**e**), and their relative abundance in coronary artery segments lacking or bearing atheroma (**f**). Cell type abundance was evaluated by chi-square test;*p<0.01, **p<0.001, and ***p<10^-8^. **g-n**, Volcano plots depicting significant differentially-expressed genes (DEGs) in atheroma versus non-atheroma-containing segments in CD8+ T cells (**g**), CD4+/CD8+ T cells (**h**), NK cells (**i**), activated NK cells (**j**), B cells (**k**), vascular smooth muscle cells (SMC, **l**), fibromyoblasts (**m**), and macrophages (**n**). Log 2 fold-change >0.2, and adjusted p<0.01. Red dots indicate genes upregulated and blue dots indicate genes downregulated in atheroma segments.

## RESULTS

### Vascular cell populations and their transcriptomes change with atheroma formation

In atheroma and non-atheroma coronary artery segments from which the adventitia was purposefully removed, thirteen cell clusters were identified (Fig. 1b,c). Cell types were assigned (Fig. 1d,e), and the most abundant populations were CD8+ T cells, CD4+/CD8+ T cells and macrophages (Fig. 1f). In atheroma-bearing segments compared to those lacking atheroma, B cells, endothelial cells, plasma cells, NK cells and fibromyoblasts were more abundant, and CD8+ T cells, macrophages, vascular smooth muscle cells (SMC) and activated NK cells were less abundant.

Volcano plots and pathway analyses reveal and categorize the differentially-expressed genes (DEGs) in the cell populations in atheroma versus no atheroma segments (Fig. 1g-n, Extended Data Fig. 1,2). CD8+ and CD4+/CD8+ T cells in atheromas display marked upregulation of metallothioneins such as MT2A and MT1X, which may regulate T cell function by influencing proliferative, oxidative, apoptotic and chemotactic responses (Fig. 1g,h, Extended Data Fig. 1a,b)(6). The T cells also have enhanced expression of complement activation genes, which may regulate their immune responses, differentiation and activation(7). Along with other RNA processing genes, the lncRNA MALAT1 is downregulated in atheroma-associated T cells, and its loss would be predicted to attenuate T cell differentiation(8). In NK cells and in activated NK cells, which are known to be atherogenic and to expand necrotic core size(9), those associated with atheromas have upregulated expression of immunoglobulin and Fc receptor signaling genes (Fig. 1i,j, Extended Data Fig. 1c,d). Atheroma-related NK cells also have decreased expression of AREG, which encodes the EGF family member amphiregulin. This may be pro-inflammatory because AREG deletion from mice results in an attenuation of resolution from inflammatory challenges(10). Regarding B cells, those associated with atheroma have upregulated expression of IGLV and IGHV genes encoding immunoglobulin lambda and heavy chain variable regions, and also increased expression of genes involved in complement activation (Fig. 1k, Extended Data Fig. 2a). Greater interrogation of B cell genes may help explain the contrasting proatherogenic and anti-atherogenic properties that have been attributed to B cells(11).

In atheroma-bearing compared to atheroma-free coronary artery segments vascular smooth muscle cell (SMC) gene profiles change dramatically (Fig. 1l, Extended Data Fig. 2b). There is a loss of contractile genes including those for myosin light chains, and upregulation of genes involved in complement activation and immunoglobulin response(Fig. 1l). With the benefit of comparing the cell populations in atheroma-bearing versus atheroma-free coronary artery segments, this provides evidence of phenotype plasticity in vascular smooth muscle cells(12) during disease development in human CAD. In atheromas fibromyoblasts also have increased expression of complement and immunoglobulin-related genes (Fig. 1m, Extended Data Fig. 2c). Interestingly, atheroma SMC and fibromyoblasts both display an upregulation in ApoE, possibly modulating lipoprotein behavior in the atheroma microenvironment.

In macrophages (Fig. 1n, Extended Data Fig. 2d) one of the most downregulated genes in atheroma is ADCYAP1, which encodes pituitary adenyl cyclase activating peptide (PACAP). ADCYAP1 global null mice have accelerated atherosclerosis, indicating that PACAP is an endogenous atheroprotective neuropeptide(13). To enhance the utility of the entire dataset and facilitate the query of genes such as macrophage ADCYAP1, a web portal has been crafted that generates UMAP plots for a gene of interest in all cell types in atheroma versus non-atheroma segments, a violin plot comparing expression of the gene in all cell types, and violin plots for the gene in any of the individual 13 cell types comparing atheroma versus non-atheroma cell populations (https://ai.swmed.edu/Xu-lab-CAD/). Interrogation of ADCYAP1 using the web portal yields UMAP plots (Extended Data Fig. 3a,b) and macrophage violin plots (Extended Data Fig. 3c) that indicate that in human coronary arteries ADCYAP1 is essentially exclusively expressed in macrophages, and that with increasing atherosclerosis severity, macrophage ADCYAP1 expression is markedly attenuated (P=4.07 e-172). With this information, further studies of ADCYAP1 in atherosclerosis should focus on its regulation and function in macrophages.

Noting marked changes in the transcriptomes of the various vascular cell types with atheroma formation, the findings may aid in the understanding of CAD risk genes. This was evaluated by determining whether 320 protein-coding genes found to be related to disease risk in CAD GWAS(4, 14) are DEGs in atheroma-associated compared to non-atheroma-associated vascular cells. Ranked based on the sum of the negative log 10 p values across cell types, the top twenty GWAS-associated risk genes are shown in the dot blot in Extended Data Fig. 4a. Findings for all detected genes are provided in Supplementary Table 2. The number one gene is PLTP, which encodes phospholipid transfer protein. PLTP is a secreted protein primarily generated by the liver, it plays a role in VLDL production and HDL biosynthesis, and it is an independent positive risk factor for CAD(15). PLTP has been detected in atherosclerotic lesions in both SMC and macrophage foam cells(16). Here it is found that PLTP expression is upregulated with atheroma formation in both SMC and macrophages, demonstrating dynamics in its expression that may be important to its paracrine and autocrine actions within vascular cells. COTL1, which encodes coactosin-like F-actin binding protein, is upregulated with atheroma formation particularly in endothelial cells. According to findings in cardiac fibrosis(17), this may be a basis for increased TGF-β signaling in endothelium during atheroma generation. Extended Data Fig. 4a also reveals that MAP4, that encodes microtubule-associated protein 4, is downregulated in a number of vascular cell types in atheroma. Its loss may cause microtubule damage and resulting cellular dysfunction(18). Furthermore, PROCR, which encodes the endothelial protein C receptor (EPCR) is downregulated in atheroma endothelium. Since EPCR binds activated protein C (aPC), facilitating aPC inactivation of other clotting factors(19), EPCR loss would be predicted to be prothrombotic. Exemplified by these findings, the inspection of the findings for a CAD risk gene in the setting of coronary artery atheroma formation may help indicate the vascular cell type in which the underpinnings of altered risk reside.

### Cell-to-cell communication changes with atheroma formation

The interrogation of human coronary artery segments with versus without atheroma provides the opportunity to reveal possible changes in cell-to-cell communication during the progression of atherosclerosis. This has been done for cytokines, growth factors, checkpoint proteins, and other ligand and receptor autocrine or paracrine interactions (Extended Data Fig. 5). Circos plots were generated using DEGs to evaluate possible gains of interaction, with ligand and/or receptor upregulated in the atheroma segment cells, and possible losses of interaction, with ligand and/or receptor downregulated in atheroma-associated cells. Query of cytokine ligands and receptors (Extended Data Fig. 5a) indicates that atheroma-associated macrophages display increased IL7 expression with possible autocrine impact and paracrine impact particularly on NK cells, plasma cells, B cells and endothelial cells. IL7 action on endothelial cells promotes the recruitment of the monocytes that become macrophages, thus promoting the inflammatory processes that underlie atherogenesis(20). In addition, in atheroma-associated macrophages, endothelial cells and SMC most cytokine receptors such as CCR1, CCR7 and CXCR4 are upregulated. Regarding loss of cytokine-related interaction, in atheroma segment CD4+/CD8+ T cells IL13 expression is downregulated with possible consequences on CD8+ T cells, SMC and NK cells, which also display a loss of the cytokine receptor CCR3. The loss of IL13 may result in a less favorable lesion morphology(21). Extended Data Fig. 5b shows the circos plot for growth factors. In atheroma both growth factor ligands and receptors are downregulated in SMC, fibromyoblasts and CD8+ T cells, with notable loss of transforming growth factor-beta 2 (TGF-β2) expression in fibromyoblasts and fibroblast growth factor (FGF) expression in endothelial cells. The loss of TGF-β2 may lead to a decrease in plaque stability(22), and the loss of FGFs may contribute to a decline in endothelial barrier function(23).

As for checkpoint ligands and receptors (Extended Data Fig. 5c), in atheroma-associated macrophages multiple checkpoint receptors are upregulated, including NFRSF14, CD28, HAVCR2, CD247, TRAF3, CD40, LTBR, ITGB2 and ITGAM. In CD4+/CD8+ T cells the checkpoint ligand CD40LG is decreased, possibly impacting CD8+ T cells and B cells in which the related receptors TRAF3, ITGB2, ITGAM and CD40 are downregulated. In atheromas B cell TNFSF4 expression is decreased, possibly impacting mechanisms in CD8+ and CD4+/CD8+ T cells in which the receptor TRAF2 is also downregulated. These cell type-specific findings may provide more granular understanding of the roles of various immune checkpoint pathways in atherogenesis, and also shed light on the association between immune checkpoint inhibitor therapy and increased CAD risk(24).

Regarding other ligand and receptor pairs possibly mediating cell-to-cell communication (Extended Data Fig. 5d), in atheromas the expression of HP, which encodes haptoglobin, in increased in CD8+ and CD8+/CD4+ T cells and fibromyoblasts. The increased haptoglobin may act via ITGAM and ITGB2 and other receptors to have autocrine and paracrine action in CD8+ T cells and paracrine actions in macrophages and B cells. The HP rs72294371 polymorphism has been linked to the incidence of CAD in Asian individuals(25). Although primarily expressed in the liver, these findings are the first to suggest that there are important paracrine mechanisms of haptoglobin action in the coronary artery. In atheroma-associated SMC, SELL (Selectin L) and KLRD1 are upregulated. Since Selectin L is an adhesion molecule primarily expressed in leukocytes and KLRD1 is an immune-activating receptor expressed in NK cells(26, 27), their upregulation in SMC may be another reflection of their capacity to switch phenotypes. Collectively, along with cell type-specific observations that are informative, these findings reveal that human coronary artery vascular cell-to-cell communication related to cytokines likely increases with greater severity of CAD, while that related to growth factors and checkpoint ligands and receptors decreases. New hypotheses related to the impact of changes in cell-to-cell communication can now be raised and tested.

### Endothelial SR-BI is upregulated in atheromas and with hypercholesterolemia

In the present study the adventitia and its accompanying vasa vasorum-associated endothelial cells have been purposefully omitted, making it possible to gain insights into mechanisms specifically occurring in the luminal blood endothelial cells. The interrogation of endothelial cell transcriptomes reveals a number of DEGs in atheroma versus non-atheroma segments (Fig. 2a). MGP, which encodes matrix Gla protein, is downregulated in atheroma endothelium, and its overexpression in mice results in decreased atherosclerotic lesion size, calcification and inflammation (28), indicating that it has anti-atherogenetic properties. In atheroma endothelium ITLN1, which encodes the adipokine intelectin-1, is the most downregulated gene. Intelectin-1 inhibits oxidative stress and reduces apoptosis, and it may have actions in endothelium mediated by Akt and eNOS signaling (29). This raises the possibility that in human atherosclerosis endothelial ITLN1 downregulation underlies a loss of the anti-atherogenic actions of NO. The metalloproteinase ADAMTS-9 is upregulated in atheroma endothelial cells. GWAS studies have revealed an association between ADAMTS-9 and the development of atherosclerosis, and serum ADAMTS-9 is increased in CAD patients compared with controls(30). Previously recognized as a microvascular endothelial cell gene(31), the now demonstrated marked upregulation of ADAMTS-9 in atheroma-associated endothelium may explain why serum ADAMTS-9 is a biomarker for CAD. The gene with the greatest relative upregulation in the endothelium associated with atheromas is the atypical chemokine receptor ACKR1. Global knockout of ACKR1 in mice results in a decrease in atherosclerotic lesion severity(32), indicating that it is pro-atherogenic. Now ACKR1 regulation and action specifically in endothelial cells warrants greater attention. Pathway analysis reveals upregulation of genes involved in inflammatory responses, which is not surprising, and in complement activation (Fig. 2b). Downregulated genes include those involved in blood vessel development.

**Fig. 2:**
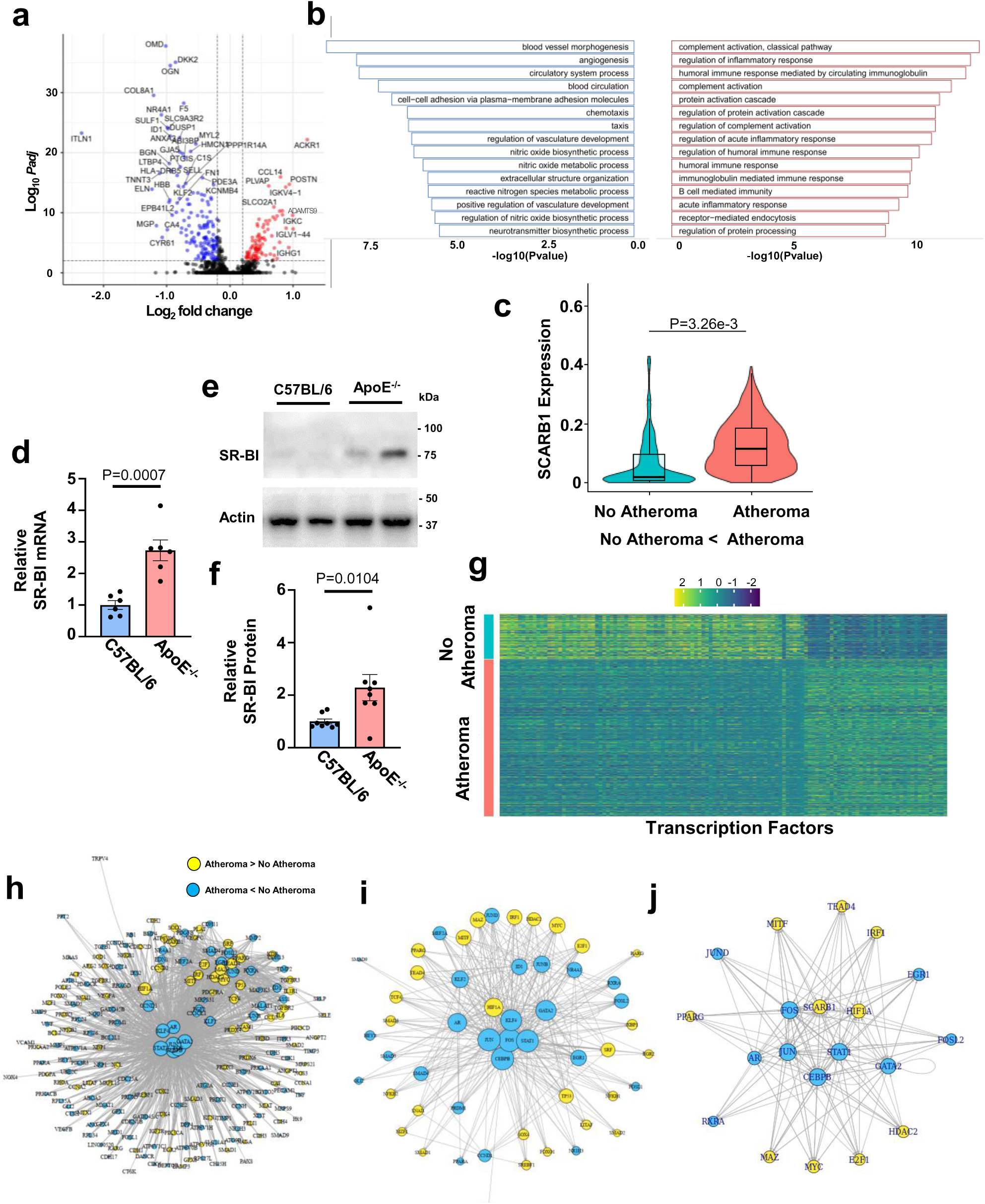
Endothelial cell SR-BI is upregulated in atheroma and with hypercholesterolemia. **a**, Volcano plot depicting significant differentially-expressed genes (DEGs) in endothelial cells, in atheroma versus non-atheroma-containing segments. Log 2 fold-change >0.2, and adjusted p<0.01. Red dots indicate genes upregulated and blue dots indicate genes downregulated in atheroma-associated endothelial cells. **b**, Top 15 pathways that are more prevalent (red) or less prevalent (blue) in endothelial cells in atheroma-containing segments compared to atheroma-free segments. **c**, Violin plots of Scarb1 expression in endothelial cells in non-atheroma versus atheroma segments. **d-f**, Effect of hypercholesterolemia on endothelial cell SR-BI mRNA abundance (**d**) and protein abundance in mouse aortic endothelial cells (**e,f**). Endothelial cells were obtained from control C57BL/6 mice on standard diet and hypercholesterolemic ApoE^-/-^mice. Representative immunoblot showing 2 samples per group is in **e**, and summary data are in **f**. **g**, Heat map of transcription factor expression in endothelial cells in non-atheroma versus atheroma segments. **h-j**, Transcription factor (TF) networks for DEGs in endothelial cells in non-atheroma versus atheroma segments. Each symbol represents a gene, those upregulated in atheroma endothelium are yellow and those downregulated are blue. Lines connecting symbols represent possible regulatory relationships between the genes, and the size of the symbol indicates the size of the node. **h**, Overall TF network including TF and target genes. **i**, Network of TF only. **j**, TF network specifically for Scarb1. In **d** and **f,** data are mean±SEM, and p values by two-tailed Student’s unpaired t text (**d**) and Mann-Whitney are shown (**f**).

With a specific mission to fill knowledge gaps about SR-BI (encoded by Scarb1) in the endothelium in atherosclerosis in humans, it was determined that SR-BI is markedly upregulated in atheroma-related endothelial cells (Fig. 2c). This prompted studies to determine if endothelial SR-BI is upregulated in the setting of hypercholesterolemia in mice. Paralleling the findings in human coronary artery endothelium, both SR-BI mRNA and protein abundance are increased in the aortic endothelium of hypercholesterolemic ApoE null mice compared to control C57BL/6 mice on standard diet (Fig. 2d-f).

Next seeking to raise possible hypotheses about how endothelial SR-BI is regulated in human atherosclerosis, it is notable that one class of endothelial cell genes particularly differentially expressed in atheroma versus non-atheroma segments is transcription factors (Fig. 2g). This suggests that the organization of endothelial cell DEGs into transcription factor (TF) networks may be informative (33, 34). Fig. 2h shows the overall TF network including TF and target genes in atheroma versus non-atheroma endothelium. Each symbol represents a gene, those upregulated in atheroma endothelium are yellow and those downregulated are blue. Lines connecting symbols represent possible regulatory relationships between the genes, and the size of the symbol indicates the size of the node. Fig. 2i shows the network for TF only, and in support of the use of TF networks for mechanistic inquiry, it reveals some predicted findings. HIF-1α, which has a major role in endothelium in atherogenesis(35), is upregulated in atheroma, it has a number of possible regulatory relationships with other transcription factors, and NF-κB1 and NF-κB2 are also upregulated. To initially interrogate processes regulating endothelial SR-BI, a TF network specifically for Scarb1 was generated (Fig. 2j), and multiple possible control mechanisms in atheroma are revealed. Thus, along with a number of other genes, endothelial SR-BI expression is altered in the setting of atheroma formation and hypercholesterolemia, and a TF network suggests multiple possible mechanisms underlying its regulation.

### Endothelial SR-BI is transcriptionally upregulated by HIF-1α

Noting that the TF network in Fig. 2j indicates parallel upregulation of HIF-1α and SR-BI mRNA in atheroma-associated endothelium in CAD, and a possible regulatory relationship between them, the impact of hypercholesterolemia on endothelial HIF-1α expression was added to the query of SR-BI in the mouse aortic endothelium. Paralleling the upregulation of endothelial SR-BI (Fig. 2d-f), HIF-1α mRNA abundance was upregulated in hypercholesterolemic mice (Fig. 3a). Recognizing that HIF-1α abundance is regulated both transcriptionally and through alterations in prolyl hydroxylation-based degradation, and by conditions other than varying oxygenation(36), the impact of HIF-1α on endothelial SR-BI expression was then interrogated in human aortic endothelial cells (HAEC) using the HIF prolyl hydroxylase inhibitor dimethyloxalylglycine (DMOG)(37). DMOG predictably caused an increase in HIF-1α protein abundance and in the expression of the known HIF-1α target VEGF (Fig. 3b,c). With DMOG, SR-BI mRNA and protein expression rose (Fig. 3d-f), and endothelial LDL transcytosis increased in an SR-BI-dependent manner (Fig. 3g,h). In HAEC the introduction of a constitutively-active HIF-1α variant had identical effects, upregulating VEGF, SR-BI mRNA and protein abundance, and LDL transcytosis (Extended Data Fig. 6). Thus, it is demonstrated for the first time that under conditions in which oxygenation is not altered, HIF-1α causes upregulation of endothelial SR-BI.

**Fig. 3:**
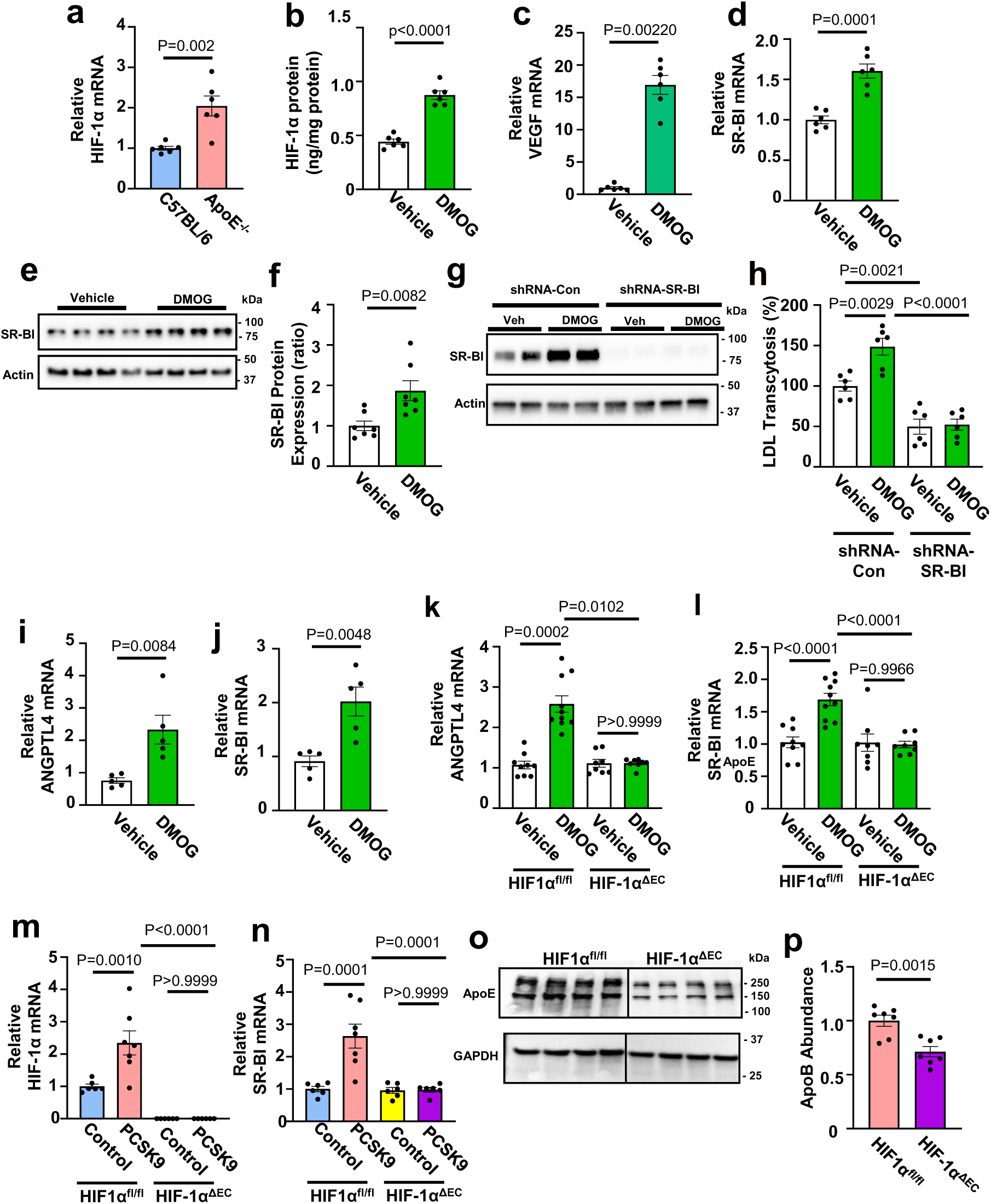
Endothelial cell SR-BI and LDL transcytosis are upregulated by hypercholesterolemia via HIF-1α. a, Effect of hypercholesterolemia on endothelial cell HIF-1α mRNA abundance in mouse aortic endothelial cells. Endothelial cells were obtained from control C57BL/6 mice on standard diet and hypercholesterolemic ApoE^-/-^ mice. **b-f**, HIF-1α increases SR-BI expression in human aortic endothelial cells (HAEC). Cells were treated with DMSO, the control vehicle, or the HIF prolyl hydroxylase inhibitor dimethyloxalylglycine (DMOG) for 24h. HIF-1α protein abundance increased (**b**) and VEGF mRNA expression rose (**c**). The DMOG treatment also causes increases in SR-BI mRNA (**d**) and protein abundance (**e,f**). Representative immunoblot showing 4 samples per group is in **e**, and summary data are in **f**. **g,h**, In HAEC the DMOG treatment increased endothelial cell LDL transcytosis in an SR-BI-dependent manner. Following control RNAi or RNAi targeting SR-BI, cells were treated with DMSO control vehicle or DMOG, and SR-BI protein abundance was evaluated by immunoblotting (**g**, 2 samples shown per group) or the transcytosis of DiI-labeled LDL was evaluated (**h**). **i,j**, DMOG treatment upregulates SR-BI in the mouse aortic endothelium in vivo. Wild-type mice were treated with DMSO vehicle control or DMOG daily for 3d, aortic endothelial cells were isolated one day later, and qPCR was performed for the known HIF-1α target gene ANGPTL4 (**i**) or for SR-BI (**j**). **k,l**, The effect of DMOG on ANGPTL4 (**k**) and SR-BI (**l**) expression was also assessed in aortic endothelium in control HIF-1α^fl/fl^ mice and in mice deficient in endothelial cell HIF-1α (HIF-1α^ΔEC^). **m-p**, The role if HIF-1α in the upregulation of endothelial SR-BI by hypercholesterolemia was tested by employing AAV8-GFP control or AAV8-PCSK9 to raise circulating cholesterol in HIF-1α^fl/fl^ and HIF-1α^ΔEC^ mice. Hypercholesterolemia increased endothelial HIF-1α and SR-BI in control mice, but not in mice lacking endothelial HIF-1α (**m, n**). **o,p**, Aorta LDL uptake, evaluated by quantifying human apoB in tissue homogenates 4h following the systemic administration of DiI-labeled nLDL, was lower in mice lacking endothelial cell HIF-1α. Representative immunoblot showing 4 samples per group is in **o**, and summary data are in **p**. Data are mean±SEM. In **a,b,d,f,i,j** and **p**, p values by two-tailed unpaired Student’s t tests are shown. Mann-Whitney was used in **c**, in **h** and **l-n** comparisons were made using One-way ANOVA with Tukey’s post-hoc testing, and p values by Kruskal-Wallis with Dunn’s post-hoc testing are shown in **k**.

To interrogate the mechanisms in vivo, DMOG was administered to wild-type mice, and it caused upregulation of known HIF-1α target genes such as ANGPTL4 and of SR-BI in the aortic endothelium (Fig. 3i,j). To then directly evaluate the actions of HIF-1α in endothelium in vivo, DMOG was administered to floxed HIF-1α control mice (HIF-1α^fl/fl^) and to mice deficient in HIF-1α selectively in endothelial cells (HIF-1α^ΔEC^). HIF-1α target genes such as ANGPTL4 were upregulated and SR-BI was upregulated by DMOG in control mice, but not in those lacking HIF-1α in endothelium (Fig. 3k,l). Thus, endothelial HIF-1α modulates SR-BI in vivo. Whether this is operative in vivo in pathogenic circumstances was then determined. Using AAV8 encoding constitutively-active PCSK9 and a hypercholesterolemic diet(1), paralleling the findings with ApoE deficiency (Fig. 2d-f, Fig. 3a), hypercholesterolemia upregulated endothelial HIF-1α and SR-BI in HIF-1α^fl/fl^ control mice (Fig. 3m,n). In contrast, hypercholesterolemia failed to upregulate SR-BI in HIF-1α^ΔEC^ mice. Importantly, in hypercholesterolemic mice endothelial HIF-1α deficiency also lowered arterial LDL uptake (Fig. 3o,p). These findings demonstrate that hypercholesterolemia causes an increase in HIF-1α in endothelial cells that leads to endothelial cell SR-BI upregulation, and a resulting promotion of LDL incorporation into the artery wall.

In the context of atherosclerosis HIF-1α is known to modulate lipid metabolism, particularly in macrophages(35). Recognizing that there are many HIF-1α target genes, and that they may mediate the influence of HIF-1α on endothelial cell SR-BI in a secondary manner, it was important to determine if the actions of HIF-1α on the receptor are direct or indirect. Guided by previously reported endothelial cell ChIP-seq data(38) and using HIF-1α binding to a regulatory element in VEGF as a positive control (Fig. 4a-c), we demonstrate in HAEC by ChIP-PCR and ChIP qPCR that HIF-1α binds to a site in Scarb1 intron 1, and that binding increases with DMOG (Fig. 4d,e). To test the consequences of HIF-1α recruitment to Scarb1 intron 1, CRISPR-Cas9 was used to delete the relevant HIF-1α binding site (Extended Data Fig. 7). Without affecting DMOG enhancement of HIF-1α abundance or VEGF expression (Fig. 4f,g), deletion of the binding site fully prevents SR-BI mRNA and protein upregulation by DMOG (Fig. 4h-j). Thus, in addition to revealing an increase in endothelial SR-BI in the setting of atheroma formation and hypercholesterolemia driven by HIF-1α that enhances artery LDL delivery, the data reveal the first direct transcriptional control of endothelial SR-BI. To facilitate inquiry of this type into the regulation of other vascular cell genes in human CAD, we have created a feature in the web portal that generates a TF network for a gene of choice in a cell type of interest from DEGs in atheroma-bearing versus non-atheroma segments (https://ai.swmed.edu/Xu-lab-CAD/).

**Fig. 4:**
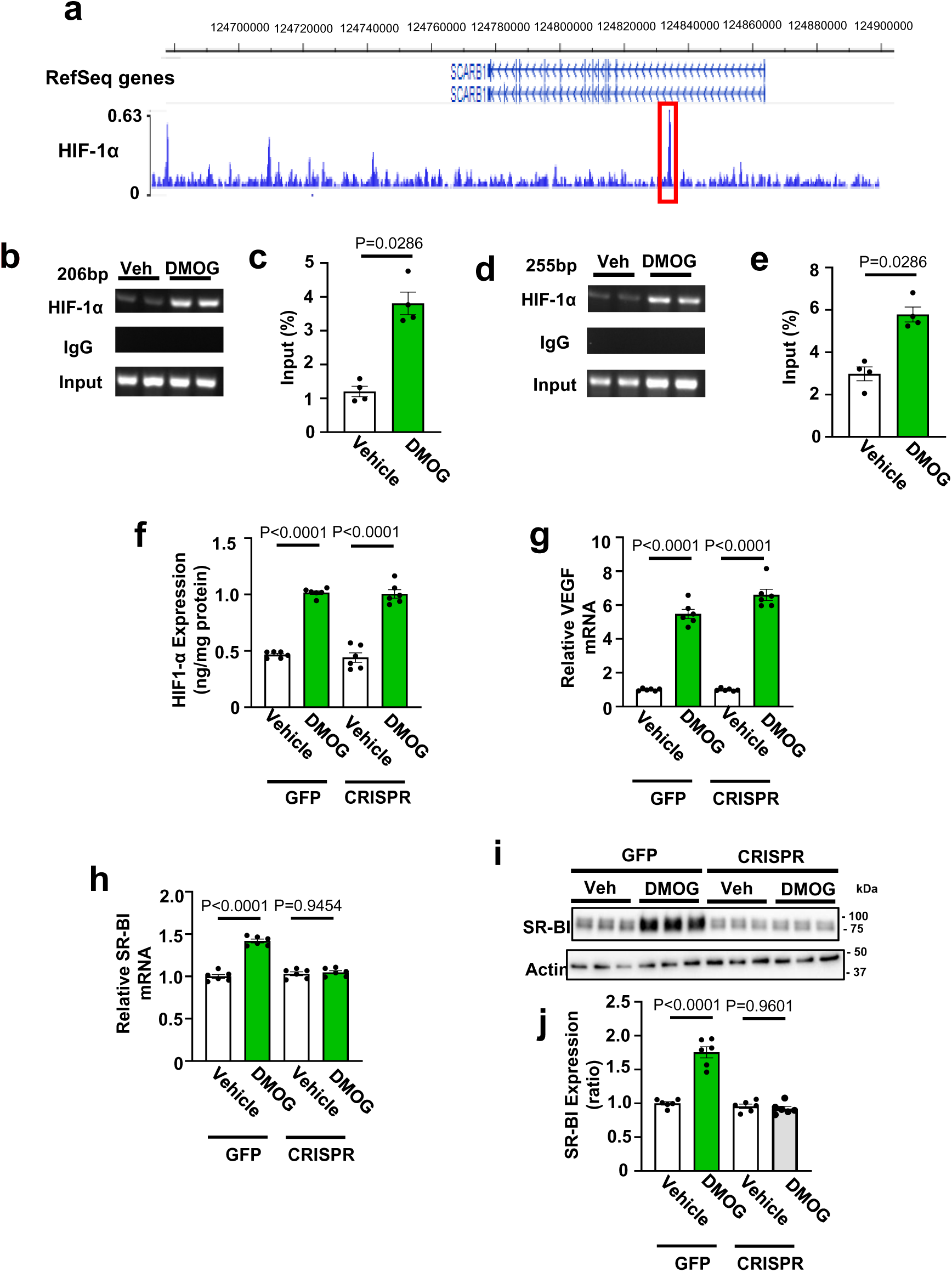
Endothelial cell SR-BI upregulation by HIF-1α is through direct transcriptional control. **a**, HIF-1α binding to intron 1 of Scarb1 revealed by ChIP-seq in hypoxia-exposed human endothelial cells (38). **b,c**, ChIP-PCR (**b**) and ChIP-qPCR (**c**) evaluating HIF-1α binding to the known HIF-1α binding site on VEGF were performed 24h following HAEC treatment with DMSO control vehicle or DMOG. **d,e**, ChIP-PCR (**d**) and ChIP-qPCR (**e**) evaluating HIF-1α binding to the predicted HIF-1α binding site on intron 1 of human Scarb1 were performed following HAEC treatment with control vehicle or DMOG. In **b** and **d** the ChIP-PCR was performed with both anti-HIF-1α antibody and an unrelated IgG control. **f-j**, The effect of DMOG on SR-BI expression in HAEC was evaluated 48h following lentiviral introduction of GFP control or CRISPR-Cas9 and two gRNA designed to delete the HIF-1α binding site in intron 1 of Scarb1. **f,** HIF-1α protein was quantified following the CRISPR manipulation and DMSO control or DMOG treatment. **g**, VEGF expression was quantified by qPCR following the CRISPR manipulation and vehicle or DMOG treatment. **h-j**, Following the same interventions SR-BI mRNA abundance (**h**) and protein abundance (**i,j**) were evaluated. Representative immunoblot showing 3 samples per group is in **i**, and summary data are in **j**. Data are mean±SEM. In **c** and **e,** p values by Mann-Whitney are shown, and in **f,g,h** and **j** the comparisons were made by One-way ANOVA with Tukey’s post-hoc testing.

### Endothelial cell subtype prevalence and gene profile change with atheroma formation

In addition to revealing new information about individual endothelial genes such as SR-BI, the interrogation of the endothelial cell transcriptomes indicates that there are three subpopulations of luminal endothelium in atherosclerotic human coronary arteries (Fig. 5a,b). The relative abundance of the endothelial cell subcluster designated as EC0 increased from 29% to 51% in the atheroma-bearing compared to the non-atheroma segments, the abundance of EC1 rose from 18 to 28%, and in contrast EC1 fell from 53 to 21% (Fig. 5c). Relative to EC1 and EC2, EC0 has a gene profile that is enriched in leukocyte activation and nucleoside metabolism genes, and deficient in genes related to the regulation of SMC proliferation (Fig. 5d.e). EC1 compared to the other subclusters is enriched in genes related to chemotaxis and the extracellular matrix (Fig. 5f,g). EC2 is enriched in genes involved in lipid handling and relatively deficient in genes related to nucleoside metabolism (Fig. 5h,i).

**Fig. 5:**
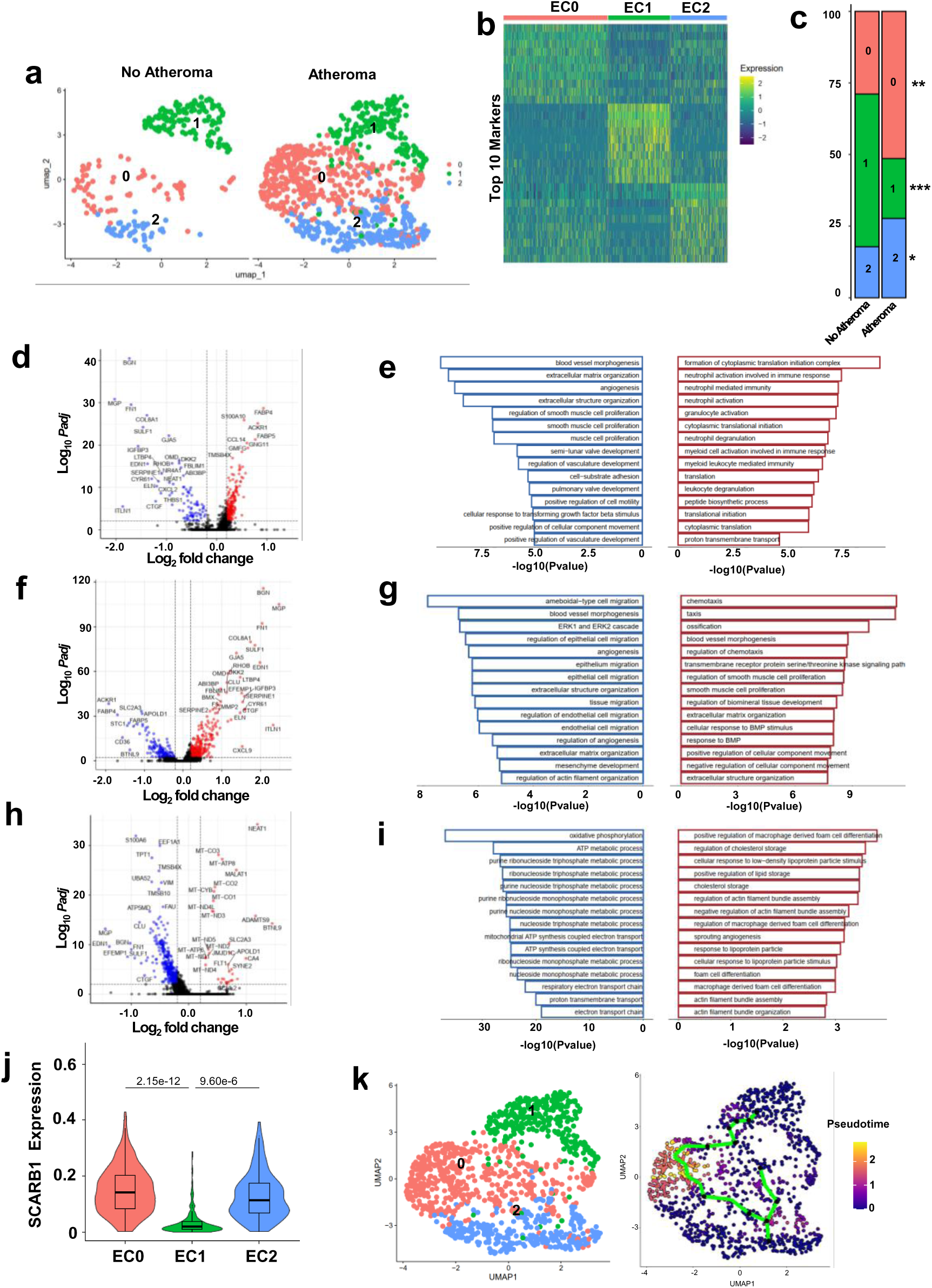
Lumenal endothelial cell subtypes and their transcriptomes change with atheroma formation. **a.** UMAP plot illustrating three endothelial cell sub sets in coronary artery segments lacking or containing atheroma. **b**, Heat map displaying top 10 most abundant genes in each endothelial cell subcluster. **c**, Distribution of endothelial cell subtypes changes with atheroma formation, with an increase in EC0 and EC2 cells and a decrease in EC1 cells. Cell type abundance was evaluated by chi-square test; *p<0.05,**p<10^-4^, ***p<10^-10^. **d**, Volcano plot depicting differentially-expressed genes (DEGs) in EC0 versus EC1 and EC2 endothelial cell subpopulations. Log 2 fold-change >0.2, and adjusted p<0.01. Red dots indicate genes upregulated and blue dots indicate genes downregulated in EC0. **e**, Top 15 pathways that are more prevalent (red) or less prevalent (blue) in EC0 versus EC1 and EC2. **f**, Volcano plot depicting DEGs in EC1 versus EC0 and EC2. **g**, Top 15 pathways that are more prevalent or less prevalent in EC1 versus EC0 and EC2. **h**, Volcano plot depicting DEGs in EC2 versus EC0 and EC1. **i**, Top 15 pathways that are more prevalent or less prevalent in EC2 versus EC0 and EC1. **j**, Violin plots comparing Scarb1 expression in EC0, EC1 and EC2. **k**, Pseudotime analysis indicating that EC0 may be derived from EC1 and EC2.

Focusing on particular genes in the EC1 subtype, which is the subpopulation that falls markedly in abundance with atheroma formation, EC1 cells are relatively enriched in BGN, which encodes biglycan. Bigycan is a small leucine-rich proteoglycan that may have atheroprotective effects. BGN knockout mice have exaggerated atherosclerosis which may be due to the loss of biglycan inhibition of thrombin activity, platelet activation and macrophage-related inflammation(39). EC1 is also enriched in MGP, which encodes matrix Gla protein which is atheroprotective, as mentioned above. Furthermore, EC1 endothelium have relatively greater expression of FN1, which encodes fibronectin 1. Plasma levels of fibronectin 1 protein are negatively associated with CAD risk, and the evidence of a cardioprotective role of fibronectin 1 is strengthened by the finding that an L16Q variant in the FN1 signal peptide which decreases fibronectin 1 secretion is associated with greater disease risk(40). Using the UMAP/Violin feature in the web portal, we find that whereas FN1 expression is observed in vascular SMC, fibromyoblasts and endothelial cells, it is in endothelial cells that FN1 expression has the most remarkable decrease in the setting of atheroma formation (Extended Data Fig. 3d-i). Thus, vascular cells, particularly the endothelium, and not the liver (41) may be the cell source responsible for the fall in plasma fibronectin 1 that is a harbinger of atherosclerosis. In addition to being enriched in matrisome proteins including bioglycan, matrix Gla protein and fibronectin 1, EC1 is the subtype of endothelial cells with the lowest expression of SR-BI (Fig. 5j). Thus, EC1 cells have numerous anti-atherogenic features, and their relative loss during atheroma formation may contribute to disease pathogenesis.

While EC1 is decreased in relative abundance in atheroma-associated coronary artery segments compared to those lacking atheroma, EC0 prevalence increases (Fig. 5a,c). To determine how the potentially more atherogenic EC0 subpopulation increases in prevalence with atherosclerosis severity, pseudotime analysis has been performed (Fig. 5k). It indicates that EC0 may arise from EC1, with an additional possible contribution from EC2.

In addition to observing changes in the prevalence of the three endothelial cell subtypes with atheroma formation, the transcriptomes within the subtypes change.

In atheroma-bearing segments, EC0 cells have increased expression of cell migration and inflammatory response genes and a loss of cytoskeletal genes (Extended Data Fig. 8a,b). EC1 endothelial cells display a loss of matrix, NO synthesis and chemotaxis genes (Extended Data Fig. 8c,d), and EC2 display an increase in vascular inflammation genes and a loss of vascular development and angiogenesis genes (Extended Data Fig. 8e,f). These collective findings suggest that in human coronary artery luminal endothelial cells there are subpopulations that change in abundance, and in their transcriptomes during atherogenesis.

### Transcript signatures can categorize endothelial cell responses to blood flow

One key mechanism governing endothelial cell transcriptomes and phenotype is the mechanosensing that occurs in response to varying blood flow characteristics. As a result of the impact on endothelial cells, unidirectional blood flow is anti-atherogenic and disturbed blood flow is pro-atherogenic(42). We leveraged the human coronary artery scRNA-seq data to determine if gene profiles in vivo in humans can identify endothelial cells which have responded to unidirectional versus disturbed blood flow. Endothelial cells were divided into those with low, intermediate or high levels of expression (bottom 1/3, middle 1/3, or top 1/3) of KLF2, which is one of the key transcription factors (TF) mediating the anti-atherogenic effects of unidirectional flow (42) (Fig. 6a,b). We then interrogated the expression of 10 other unidirectional flow-responsive genes and found that 9 of the 10, including KLF4, display the same pattern of expression as KLF2 (Fig. 6c). This indicates that the KLF2 high (top 1/3) versus KLF2 low (bottom 1/3) endothelial cells represent those in the human coronary artery with gene signatures indicative of responding to unidirectional versus disturbed blood flow, respectively.

**Fig. 6:**
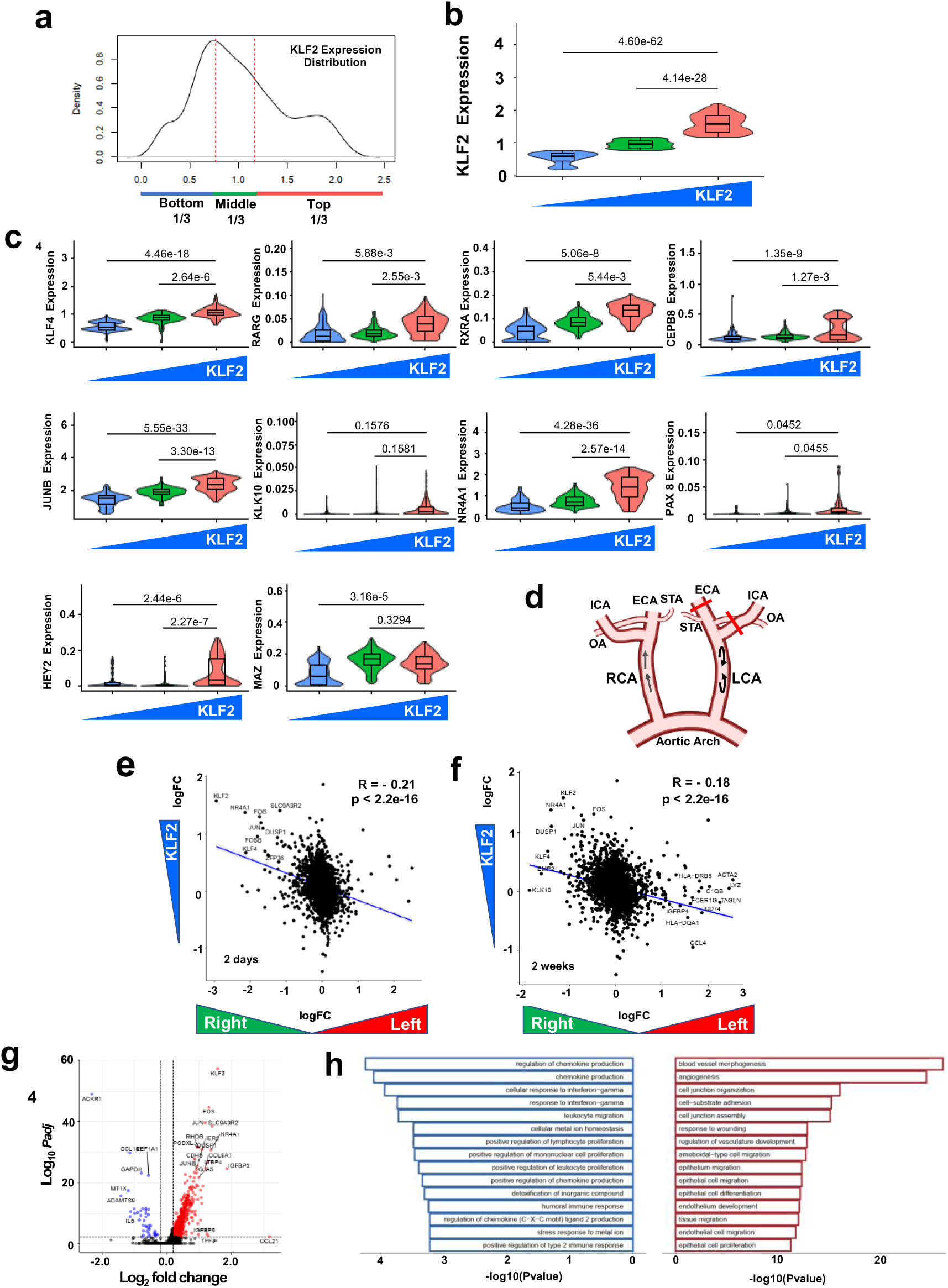
Transcript signatures can categorize endothelial cell responses to blood are divided into those with low, intermediate or high levels of KLF2 expression (bottom 1/3, middle 1/3, or top 1/3). **b**, Violin plots depicting the relative levels of expression of KLF2 in the three abundance categories. **c**, Violin plots depicting the relative levels of expression of ten other unidirectional flow-responsive genes related to the spectrum of KLF2 expression. **d**, Schematic of the mouse carotid artery partial ligation model of disturbed blood flow. The ligation of major branches of the left carotid artery (LCA) results in disturbed flow and the right carotid artery (RCA) provides the control condition of linear, unidirectional flow. Three of the four caudal branches of the LCA, the left external carotid artery (ECA), internal carotid artery (ICA), and occipital artery (OA) are ligated and the superior thyroid artery (STA) is left intact. **e,f**, Correlation between the differentially expressed genes in KLF2 low (bottom 1/3) versus KLF2 high (top 1/3) human coronary artery endothelial cells and the murine endothelial cell genes with differential expression in the LCA versus RCA endothelium 2 days (**e**) or 2 weeks (**f**) post-ligation. **g,** Volcano plot depicting differentially-expressed genes (DEGs) in KLF2 high compared to KLF2 low human coronary artery endothelium. Log 2 fold-change >0.2, and adjusted p<0.01. Red dots indicate genes upregulated in KLF2 high and blue dots indicate genes downregulated in KLF2 high. **h**, Top 15 pathways that are more prevalent (red) or less prevalent (blue) in KLF2 high compared to KLF2 low endothelial cells.

To evaluate the observations made employing the gene profiles related to flow responses, the human coronary artery endothelial cell findings for KLF2 high versus KLF2 low have been compared to publically-available single cell RNAseq data for intimal cells in the mouse carotid artery partial ligation model of disturbed blood flow (Fig. 6d). The ligation of major branches of the left carotid artery (LCA) results in disturbed flow and the right carotid artery (RCA) provides the control condition of linear, unidirectional flow(43, 44). In Fig. 6e there is a correlation between the differentially expressed genes in KLF2 low versus KLF2 high human coronary artery endothelial cells and the murine endothelial cell genes with differential expression in the left versus right carotid artery endothelium two days post-ligation (p<2.2 e-16). There is also a correlation with the differentially expressed murine genes two weeks post-ligation (Fig. 6f, p<2.2 e-16). For example, NR4A1, an orphan nuclear receptor that promotes angiogenesis(45) and DUSP1, a negative regulator of adhesion molecule expression(46), are more highly expressed in parallel in the mouse right carotid artery endothelium exposed to unidirectional flow and in the human coronary endothelium with a related KLF-2 high gene signature. Two weeks post-ligation TAGLN, which encodes the actin-crosslinking protein transgelin and is expressed in the process of endothelial mesenchymal transition (47), is more abundant in the left carotid artery exposed to disturbed flow and the coronary endothelium with a parallel KLF-2 low gene profile. CD74, a binding protein for MIF which is upregulated to increase ROS and NF-kB in injured endothelium(48), is also increased in parallel in the human and mouse endothelium subjected to disturbed blood flow. A similar observation is made for CCL4, which encodes a chemokine that activates inflammatory signaling and alters the distribution of junctional proteins(49). Although this comparison is between human coronary artery endothelium and murine carotid artery endothelium, it provides evidence that the KLF2 high versus KLF2 low gene profiles have efficacy revealing how endothelial cells respond to varying flow in vivo in the human coronary artery.

Having evidence of utility of the KLF2-based categorization of human endothelial cell responses to blood flow, DEGs were evaluated (Fig. 6g). One of the most downregulated genes in KLF2 high endothelium, indicative of greater expression in the setting of disturbed flow, is ADAMTS-9, and it is mentioned above that ADAMTS-9 is increased in coronary artery disease (CAD) patients compared with controls(30). ADAMTS-9 is anti-angiogenic and it acts cell-autonomously in endothelial cells(31), and we now show that it is likely upregulated by disturbed blood flow in the human coronary artery. Above we mentioned that ACKR1, or atypical chemokine receptor-1, is highly expressed in the endothelium of atheroma-bearing coronary artery segments (Fig. 2a), and here we find that it is decreased in KLF2 high endothelium, suggesting that it is upregulated by disturbed blood flow. It is also notable that MT1X, which is a metallothionein upregulated in endothelial cells by hypoxia(50), is upregulated in the human coronary artery in endothelium with a gene profile indicative of response to relatively disturbed blood flow. Another gene downregulated in KLF2 high endothelial cells is CCL14, encoding C-C Motif Chemokine Ligand 14, which is found in greater abundance in vulnerable compared to stable carotid artery plaques(51). This suggests that its expression is enhanced in endothelium in regions of disturbed blood flow, possibly contributing to the increase in plaque instability in those locations(52).

Regarding genes more highly expressed in KLF2 high endothelium, they include PODXL, which encodes podocalyxin. Podocalyxin is a component of the glycocalyx on the luminal face of most blood vessels, and it attenuates leukocyte-endothelial cell adhesion and maintains endothelial barrier function, particularly under inflammatory conditions(53, 54). Another gene now found to likely be responsive to unidirectional flow is IGFBP3, which promotes vascular repair by increasing eNOS expression(55). In broader terms, pathway analysis (Fig. 6h) indicates that endothelial genes relevant to angiogenesis, cell junction assembly and response to wounding are upregulated, and genes relevant to chemokine and cytokine production and responses to IFNG are downregulated in response to unidirectional blood flow in the human coronary artery. In this manner, the use of the KLF2-based gene profile provides insight into a critical mode of endothelial cell perturbation in human arteries such as the coronary artery.

How varying blood flow characteristics influence endothelial cell genes related to CAD risk is considered in Extended Data Figure 4b. The top twenty genes ranked based on the product of the negative log 10 p values for DEGs related to flow and to atheroma formation are shown. Findings for all detected genes are provided in Supplementary Table 3. The number one gene is F5 that encodes coagulation factor V. Factor V acts as a cofactor for factor Xa and thereby has procoagulant function, but as a cofactor for aPC it has important anticoagulant function. The latter is particularly apparent in the setting of the factor V Leiden mutation (FVL), which is the most prevalent hereditary risk factor for thromboembolism(56). Using plasma and endothelial colony-forming cells from FVL carriers and controls in crossover ex vivo experiments, it has recently been revealed that endothelial-expressed factor V is the major driver of its anticoagulant function(57). With that in mind, the current data indicating that its expression in endothelial cells is upregulated by unidirectional blood flow and dowregulated in atheromas suggests for the first time that the loss of endothelial cell factor V may play a role in coronary artery atherothrombosis. Extended Data Figure 4b also indicates that PI16 in endothelium is upregulated in the setting of unidirectional blood flow and downregulated in atheroma. Encoding peptidase inhibitor 16 protein, which is a shear stress- and inflammation-regulated inhibitor of MMP2 that supports the maintenance of endothelial cell quiescence(58), its loss with disturbed blood flow and in atheroma may detrimentally impact endothelial cell phenotype. The consideration of endothelial cell DEGs related to flow and atheroma formation in the current work may provide insights into the basis for the impact of certain genes on CAD risk. To best leverage the new information now provided related to endothelial cell responses to flow, a tool has been generated in the web portal that makes it possible to use the flow-related gene signature to determine if an endothelial cell gene of interest is responsive to differential forms of flow (https://ai.swmed.edu/Xu-lab-CAD/). There is also a web portal feature that generates TF networks that reveal possible modes of regulation of the gene in response to blood flow.

### Endothelial cell SR-BI and LDL transport are upregulated by disturbed blood flow

Seeking to further understand the mechanisms regulating endothelial SR-BI in the human coronary artery, we interrogated the levels of SR-BI expression in endothelial cells with differing levels of KLF2 expression. We found that SR-BI expression is least in endothelial cells with high KLF2 compared to middle range KLF2 or low KLF2 (Fig. 7a). This prompted a query of the scRNA-seq data on endothelium in the mouse carotid artery model of disturbed (versus unidirectional) blood flow(43, 44). Paralleling the findings with KLF2 categorization of human coronary artery endothelium, by single cell RNAseq SR-BI expression in mouse intimal cells was greater in the ligated left carotid artery (LCA; disturbed flow) than in the control right carotid artery (RCA; unidirectional flow), both 2 days and 2 weeks post ligation (Fig. 7b,c). These collective findings by RNAseq suggest that SR-BI is increased by disturbed flow in vivo in both human and mouse endothelial cells. To directly determine the effect of disturbed blood flow on endothelial SR-BI expression, qPCR was performed on intimal cells in the mouse carotid artery partial ligation model. It revealed upregulation of SR-BI in the intimal cells at 2 days post-ligation, whereas KLF2 expression predictably fell (Fig. 7d). This observation raises the possibility that there is not only an association between regions of disturbed blood flow and lipid accumulation in atherosclerotic lesions(59), but that blood flow actively influences LDL transport into the artery wall. With minor modifications in our approach to image DiI-nLDL uptake in the mouse aorta(1), we established a method to do the same in en face preparations of the carotid artery. We then used confocal immunofluorescence to image nLDL uptake in the partial ligation model 2 days post-ligation in ApoE^-/-^ mice fed a hypercholesterolemic diet. Immunofluorescent labeling of VE-cadherin was employed to identify and visualize the endothelial monolayer during confocal imaging. Vehicle alone yielded no background signal for DiI, and in the non-ligated control right carotid artery (RCA) there was no DiI-LDL uptake (Fig. 7e,f). In marked contrast, DiI-LDL uptake was greatly induced by disturbed blood flow in the ligated left carotid artery (LCA). However, the induction of endothelial LDL uptake by disturbed blood flow was prevented by the deletion of SR-BI selectively in endothelial cells (Fig. 7g,h). Therefore, whereas prior studies have provided correlations between regions of disturbed flow and increased lipid accumulation(60), we show for the first time that there is a causal relationship, with disturbed blood flow robustly promoting endothelial LDL uptake. We further demonstrate that this is due to a disturbed flow-induced upregulation of endothelial SR-BI. Revealing the consequences of this newly-identified process, the deletion of endothelial SR-BI also greatly attenuated atherosclerotic lesion formation evaluated 7 days post-ligation (Fig. 7i,j). Prompted by the discoveries in the human coronary artery, these observations reveal mechanistic linkage between disturbed blood flow, endothelial SR-BI upregulation, a resulting increase in endothelial LDL transport, and disturbed flow-induced atherogenesis.

**Fig. 7:**
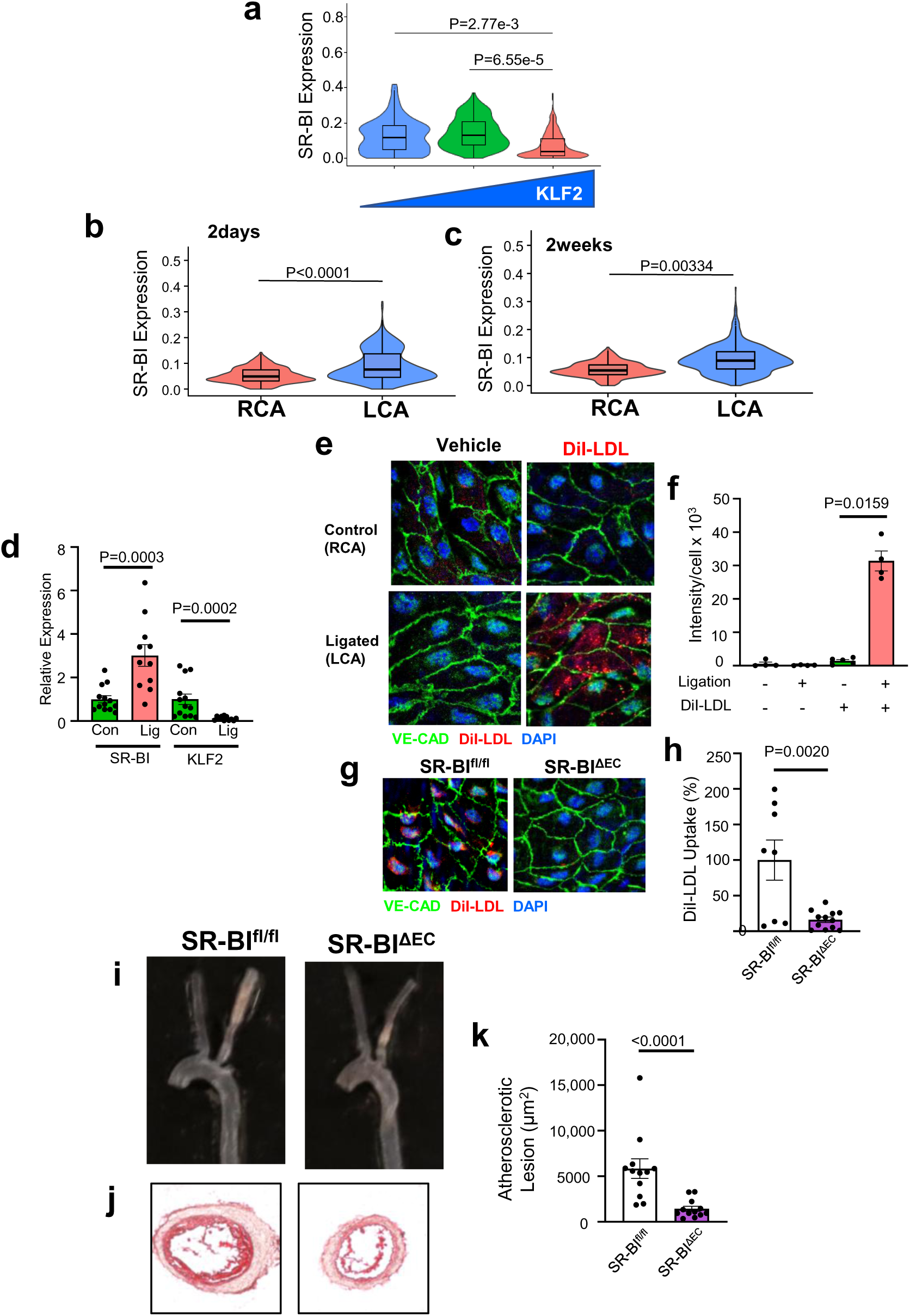
Endothelial cell SR-BI and LDL transport are upregulated by disturbed blood flow. **a**, Violin plots show relative levels of expression of S-BI in endothelial cells with KLF2 low, middle range, or high abundance of KLF2. **b,c**, Violin plots depicting relative endothelial cell SR-BI expression in the control right carotid artery (RCA) and ligated left coronary artery (LCA) in the mouse model of disturbed blood flow. The findings are from query of a publicly-available, published single cell RNAseq dataset (46), with samples obtained 2d and 2wk post-ligation. **d,** Intimal cell SR-BI expression and KLF2 expression determined by qPCR in the control RCA and ligated LCA 2d post-ligation in wild-type mice. **e**, En face images of RCA and LCA endothelium 4h following IV-administered vehicle versus DiI-LDL 2d post-ligation in ApoE^-/-^ mice. **f**, Summary data for fluorescence intensity for multiple mice studied as in **e**. **g**, En face images of LCA endothelium in SR-BI^fl/fl^ or SR-BI^ΔEC^ mice on ApoE^-/-^ background 4h following IV administration of DiI-LDL 2d post-ligation. **h**, Summary data for fluorescence intensity expressed relative to intensity in SR-BI^fl/fl^ mice, for multiple mice studied as in **g**. **i-k**, Atherosclerosis in the LCA 7d post-ligation in SR-BI^fl/fl^ or SR-BI^ΔEC^ mice on ApoE^-/-^ background. Gross images of the aortic arch, RCA and LCA (**i**) and Oil Red O-stained cross-sections (**j**) are shown. **k**, Atherosclerotic lesion size on cross-sections. In **d,f,h** and **k**, data are mean±SEM. In **d,f** and **k,** p values by Mann-Whitney are shown, and in **h** the comparison was made by two-tailed unpaired Student’s t test.

## DISCUSSION

In the present work scRNA-seq data from human coronary artery segments containing versus lacking an overt atheroma has been leveraged to provide new knowledge about the molecular mechanisms occurring in vascular cells during the progression of CAD. The removal of the adventitia and the related vasa vasora made it possible to query genes specifically in the luminal endothelium. Vast differences in transcriptomes were observed in both non-endothelial cells and endothelial cells in segments with versus without atheroma, and a web portal has been constructed to facilitate mechanistic interrogation. Recognizing the critical role of endothelial SR-BI in atherogenesis(1), we reveal that in CAD endothelial SR-BI expression is increased in the setting of atheroma formation, and in endothelial cells with a transcript signature indicative of responding to disturbed blood flow. In mice we show that independently hypercholesterolemia and disturbed blood flow upregulate endothelial SR-BI. With disturbed blood flow the SR-BI upregulation underlies the initiation of endothelial cell LDL uptake leading to atherogenesis. Guided by transcription factor networks, mechanistic studies show that HIF-1α binding to Scarb1 intron 1 governs endothelial SR-BI transcription, leading to increased LDL transcytosis. We then determine in vivo that artery LDL uptake in the setting of hypercholesterolemia is driven by HIF-1α upregulation of endothelial SR-BI. Thus, the two major instigators of atherosclerotic lesion formation, hypercholesterolemia and disturbed blood flow, both upregulate endothelial SR-BI to drive the LDL transport that underlies the disorder. The processes that regulate endothelial SR-BI may be new therapeutic targets in the battle against CAD.

The present scRNA-seq findings in CAD segments of differing atherosclerosis severity can be combined with the single-cell ATAC-seq observations of Turner et al(5) to further reveal the driver TF regulating disease-relevant genes in different cell types. The present observations related to SMC and fibromyoblasts may add to our understanding of stable versus unstable atherosclerotic plaques, such as the senescence-like versus osteogenic-like phenotype previously observed by Alloza and colleagues in asymptomatic versus symptomatic human carotid artery plaques(61). The present work may also shed light on the role of TCF21 and other genes in SMC phenotypic modulation to fibromyoblasts, which was revealed in diseased human coronary arteries by Wirka et al(62). The present findings for immune cells complement what has been revealed by Fernandez et al in clinically symptomatic versus asymptomatic carotid artery plaques(63). In asymptomatic plaques they observed that T cells and macrophages were activated by IL-1β signaling, and our web portal reveals that IL-1β is expressed primarily in macrophages, with greater levels of expression in macrophages from atheroma-bearing coronary artery segments. In the query of endothelial cell DEGs we demonstrate an upregulation of genes involved in complement activation and inflammatory response in endothelium associated with atheroma, and a decline in the expression of genes related to vascular development and nitric oxide production. In addition to being informative about changes in individual genes in specific vascular cell types, the DEG data go beyond the available cell-to-cell communication information in carotid artery endarterectomy samples(4, 63) and normal coronary arteries(64); they reveal changes in cell-to-cell cross-talk which likely occur with increasing degree of atherosclerosis in the diseased coronary artery.

With a focus on luminal blood endothelial cells, we find that there are three subpopulations in human CAD, and their prevalence and transcriptomes change with the severity of atherosclerosis. One subpopulation is enriched in leukocyte activation and nucleoside metabolism genes, and relatively deficient in genes related to SMC proliferation; it increases in abundance and has further upregulation of inflammatory genes with atheroma formation. Another subpopulation also increases in abundance and has greater vascular inflammation genes and fewer angiogenesis genes with increasing atherosclerosis severity. The third subpopulation is enriched in genes encoding ECM proteins and the atheroprotective proteins biglycan and matrix Gla protein, and it is the subpopulation with the lowest expression of SR-BI; it decreases in prevalence with atheroma formation. Thus, there are both increases in endothelial cell subpopulations with pro-atherogenic features and a decline in relatively atheroprotective endothelial cells with worsening atherosclerosis severity. These observations add new insights to those gained about endothelial cell subpopulations in prior studies related to normal vasculature or atherosclerosis. In addition to lymphatic endothelium, a prior study of normal mouse aorta found two major endothelial cell subpopulations(65). One is enriched in ECM and integrin cell surface interaction genes, and one is enriched in lipid transport and angiogenesis genes. With Western diet feeding there were similar changes in gene expression in the two subpopulations, with an increase in contractile function genes including MYL9, TAGLN and ACTA2. Interestingly, we find that in human CAD all three of these genes are decreased in expression in endothelium with atheroma formation. In a report of endothelial cell heterogeneity in the coronary artery branches of mice there were three subclusters. They were enriched in immune response, angiogenesis-related, or endothelial cell growth and vascular tone modulation genes (66). A pioneering study in human carotid artery atherosclerosis found four endothelial cell subclusters(4). One likely represents the endothelium in the vasa vasorum, two others are activated cells, and a fourth had features suggesting that the cells were undergoing endothelial-to-mesenchymal transition. An earlier study of disease-free human arteries revealed three endothelial cell subpopulations(64). Interestingly, the most prevalent subgroup in the coronary artery, comprising 74% of the endothelial cells, was enriched in inflammatory genes including ACKR1, CCL14 and SELE. Our study and these prior efforts have consistently revealed a limited number of discrete endothelial cell subpopulations. Since there is increased endothelial cell turnover in atherosclerosis-prone areas, and neighboring cells can be major contributors to endothelial cell replacement(67), in atherosclerosis one endothelial cell subpopulation may arise from another. We tested this possibility in the present dataset by performing pseudotime analysis. It revealed that the subpopulation that is enriched with proinflammatory genes, which increases to be the most abundant in coronary artery segments with atheroma, is likely derived from one or both of the other subpopulations. Lineage tracing approaches will be required in mice to reveal the sources of different endothelial cell subpopulations in the atherosclerotic artery.

It is well recognized that atherosclerotic lesions preferentially develop at vascular sites where blood flow is disturbed, including near arterial curves and branch points. The endothelium is exquisitely sensitive to differences in blood flow characteristics, with mechanosensing having a major impact on endothelial cell gene expression and phenotype. Extensive work has elucidated the underpinnings of the endothelial mechanosensing and its downstream consequences in cell culture and in animal models(68). In contrast, our understanding of how varying flow patterns regulate endothelial cell expression in vivo in humans remains limited. To bridge this knowledge gap we segregated the human coronary artery endothelial cells based on their relative expression of KLF2. Parallel relative expression of a number of flow-responsive genes, and a comparison with intimal cell transcriptomes in the mouse model of disturbed flow suggest that the KLF2-based gene signature may have utility revealing how endothelial cells respond to varying flow in vivo in the human coronary artery. The evaluation of DEGs between KLF2 high versus KLF2 low endothelial cells has revealed specific genes and functional groups of genes that may be influenced by flow.

With an initial impetus to better understand the biology of endothelial cell SR-BI in CAD, we queried whether its gene SCARB1 is differentially expressed in the endothelium of atheroma versus non-atheroma coronary artery segments. Increased transcript abundance was found in atheroma endothelium, leading to studies in mice that demonstrated that hypercholesterolemia upregulates endothelial SR-BI. Leveraging the developed gene signature informative about coronary artery endothelial cell responses to blood flow, it was determined that SR-BI is upregulated in the setting of disturbed flow compared to unidirectional flow. This prompted studies in the mouse carotid artery partial ligation model, which demonstrated that SR-BI is indeed upregulated by disturbed blood flow. Disturbed flow also caused a striking increase in endothelial cell LDL uptake which was dependent on the flow-induced SR-BI upregulation. Upregulation of the scavenger receptor was also responsible for disturbed flow induction of atherosclerosis. For many years it has been observed that lipid accumulation is increased in regions of disturbed blood flow that are prone to atherosclerotic lesion development(59). However, it has not been previously demonstrated that the disturbed blood flow drives LDL uptake, and the underpinnings of the process were not known. Whereas cell culture studies have shown that endothelial SR-BI is upregulated by HDL from healthy human donors and downregulated by HDL from diabetics and by estrogen(69, 70), the present work is the first to provide insights into endothelial SR-BI regulation in vivo.

With the availability of data on DEGs in endothelium at different degrees of atherosclerosis severity, it was possible to create TF networks to reveal possible modes of gene regulation in that setting. A TF network centered on SR-BI (SCARB1) revealed parallel upregulation of SCARB1 and HIF-1α and a possible regulatory relationship in atheroma compared to non-atheroma coronary artery endothelium. Mouse studies revealed similar parallel upregulation in aorta endothelium in the setting of hypercholesterolemia, prompting studies of HIF-1α and SR-BI in primary human aortic endothelial cells (HAEC). Increased HIF-1α stabilization with the prolyl hydroxylase inhibitor DMOG caused upregulation of SR-BI, and LDL transcytosis was increased in parallel in an SR-BI dependent manner. The underlying mechanism involved HIF-1α binding to a regulatory element in human SCARB1 intron 1, revealing the first direct transcriptional control of endothelial SR-BI. The actions of HIF-1α in atherosclerosis are complex, with evidence of both anti- and pro-atherogenic roles. In atherosclerosis there are multiple mechanisms that increase HIF-1α activation. Local hypoxic conditions stabilize HIF-1α, there are inflammatory conditions that upregulate the transcriptional control of HIF-1α, which is an NF-kB target gene, and cholesterol and oxysterols increase HIF-1α signaling (35). The new insights into HIF-1α action on Scarb1, which enacts a major mechanism underlying atherogenesis, may help reveal specific processes that possibly can be leveraged for atheroprotection.

In the present work, how the data obtained was leveraged to learn about endothelial SR-BI in atherosclerosis exemplifies how new insights can be gained about thousands of other genes in more than a dozen vascular cell types. To facilitate such queries, we have generated user-friendly features in a web portal. For a gene of interest UMAP plots and violin plots can be produced to display the cell type specificity of expression. Differences in relative expression in a cell type of interest in atheroma-bearing versus atheroma-free coronary artery segments viewed in violin plots indicate how the abundance of the transcript changes with the advancement of atherosclerotic severity. For an endothelial cell gene, violin plots can be generated related to the cell signature for blood flow response to assess how unidirectional versus disturbed flow impacts its expression in the human coronary artery. And employing DEGs related to either atherosclerosis severity or the endothelial cell flow response signature, transcription factor networks can be obtained to raise hypotheses about gene regulation. To aid in the extrapolation of the data to the genetics of CAD, changes in gene expression with atheroma formation or with differential flow conditions (endothelial cells) have been compiled for over 300 CAD risk genes. The inspection of the findings for a CAD risk gene in the setting of coronary artery atheroma formation may help indicate the vascular cell type in which the underpinnings of altered risk should be pursued. In addition, the basis for the impact of an endothelial cell gene on CAD risk may be revealed by considering if it is a DEG related to disturbed blood flow or atheroma formation. These measures should greatly extend the utility of the data set to interrogations of numerous pathogenetic mechanisms in CAD.

Valuable prior scRNA-seq and ATAC-seq studies in atherosclerotic carotid and coronary arteries have revealed the epigenetic landscape and gene expression present in vascular cells in atherosclerosis in humans(4, 5). The present work provides new insights into the dynamics of gene expression and the underlying regulatory processes operative during the progression of atherosclerosis in humans. It further reveals that independently, the two major instigators of atherosclerotic lesion formation, hypercholesterolemia and disturbed blood flow, upregulate endothelial SR-BI to drive the LDL transport that underlies the disorder. The former discovery sheds new light on how hypercholesterolemia increases artery LDL uptake(71), revealing that direct HIF-1α upregulation of endothelial SR-BI is the missing link. The latter demonstration of a flow-responsive endothelial mechanism counters the prior thinking that small gaps between endothelial cells or increased residence time of circulating LDL underlie how disturbed blood flow promotes LDL entry to the subendothelial space(60). The modulation of endothelial SR-BI expression and mechanism of action may represent new opportunities to combat the considerable residual risk of CAD that exists despite lipid lowering strategies.

### ONLINE METHODS

#### Human Specimens and Patient Clinical Parameters

Coronary arteries were obtained from hearts excised from five patients with atherosclerosis and ischemic heart disease (n=4) or non-ischemic cardiomyopathy (n=1) requiring heart transplant at UT Southwestern Medical Center. Patient clinical parameters are provided in Supplementary Table 1. The subjects gave written informed consent, and the study was approved by the Institutional Review Board at the University of Texas Southwestern Medical Center.

#### Sample Collection and Preparation and Characteristics

Coronary arteries were immediately dissected from the excised hearts of the transplant recipients. Segments were selected from the right coronary artery or the left anterior descending coronary artery. From each subject one segment was chosen that contained an obvious atheroma by visual inspection and palpation, and a second segment was chosen that lacked an atheroma. The segments were placed in ice cold normal saline until processed. Within 2h post-dissection the adventitia was removed from the segments by fine dissection, and an aliquot was removed for histological evaluation. Hematoxylin and eosin staining revealed that all the segments had plaque classifications of V or VI(72–74).

The remaining segments were cut open longitudinally to expose the intimal surface, and they were placed in digestion buffer (1x HBSS, 1%BSA, 1mM EDTA) containing the enzymatic dissociation reagents (10.4U/ml Liberase (Sigma), 1mM CaCl_2_, 0.58mg DNase I and 0.99mg Hyaluronidase (Sigma) in 10ml digestion buffer), and cut into small pieces with fine scissors. After incubation at 37°C for 1h with periodic agitation, the cell suspension was passed through a 70um cell strainer, washed twice in digestion buffer, and centrifuged at 300g for 5min. Erythrocytes were removed using red blood cell lysis buffer (Sigma). Dead cells were removed using Annexin V staining (Stemcell) according to the manufacturer’s instructions, and cells were counted with an automatic cell counter.

#### 10X Genomics Single Cell RNA-seq

Single cells were encapsulated in droplets through the GemCode single cell platform using the GemCode GelBead, chip and library kits according to the manufacturer’s instructions (1000014, 10X Genomics) to target 10,000 cells per sample. Briefly, the single cell suspension was loaded onto a well on a 10X Chromium single cell instrument, and then partitioned into nanoliter-scale Gel Bead-in-Emulsions (GEMs), in which all the cDNA generated from an individual cell shares a common 10X Barcode, followed by amplification, shearing and 5’ adaptor and sample index attachment. Qualitative analysis was performed using the Agilent Bioanalyzer High Sensitivity assay. Libraries were sequenced on a NovaSeq. Whereas two samples were sequenced separately, the rest of the 7 samples were sequenced at the same time to minimize batch differences.

#### Preprocessing of Single Cell RNA-seq Data

The Cell Ranger Software Suit (version 3.1.0) was used to perform sample demultiplexing, barcode processing and single-cell 3’ gene counting. The sequencing reads were aligned to the GRChg38 reference genome and raw count matrix were generated. The SoupX R package (version 1.5.2) was used for potential ambient RNA removal. To exclude low-quality cells, Scrublet software (version 0.2.3) was first used to identify and filter potential doublets. Further quality filtering was performed using the Seurat R package (version 3.2.2). To obtain high quality cells for downstream analysis, cells were filtered out that had < 200 detected genes or >10,000 nCount_RNA or >15% mitochondrial reads, further removing cells that could be potentially duplets or dead. After these stringent filtering approaches 5417 ‘No Atheroma’ cells and 14938 ‘Atheroma’ cells were obtained.

#### Integrative Analysis of Single Cell RNAseq Data

To enable comparison of samples from different conditions, integrated analysis of the data from all samples was performed following the published SCTransform integration procedure (https://satijalab.org/seurat/archive/v3.1/integration.html)(75). In brief, the SCTransform() function was first used to normalize each sample, then the SelecIntegrationFeatures() function was used to select features for integration and the PrepSCTIntegration() function was used to calculate necessary Pearson residuals. The FindIntegrationAnchors() function was then employed to identify anchors and the IntegrateData() function was finally used to integrate all sample datasets. The integrated data was scaled and normalized using the ScaleData() function and NormalizeData() function. RunPCA() and RunUMAP() functions were used with the first 30 principle components to perform dimensional reduction. The FindNeighbors() function and FindClusters() function were performed to accomplish unsupervised clustering, and 13 clusters resulted. The FindAllMarkers() function was used to calculate specific markers for each cluster to help with cell-type annotation. Cell types for each cluster were manually annotated by inspecting known cell type-specific gene marker expression. Differential expressed genes (DEGs) between conditions were identified using the FindMarkers() function, with p-values calculated using default Wilcoxon Rank Sum tests and adjusted p-values calculated with Bonferroni correction. To interrogate how the findings relate to CAD risk genes, it was determined whether 320 protein-coding genes found to be related to disease risk in CAD GWAS(4, 14) are DEGs in atheroma-associated compared to non-atheroma-associated vascular cells, or DEGs in KLF2 high versus KLF2 low endothelial cells.

#### Gene Ontology term Enrichment Analysis

Enrichment analysis of GO terms was performed using clusterProfiler R package (version 4.14.6), with upregulated or downregulated list of DEGs as input.

#### Cell-cell Communication Analysis

To analyze cell-cell communication changes among different cell-type pairs, we adopted the differential combination-based tool iTALK R package(76). In brief, significantly differentially expressed genes (DEGs) of each cell-type between conditions were identified using the DEG() function within the iTALK package and the ‘wilcox’ statistics method, and DEGs between two conditions were then used as input for the FindLR() function in the iTALK package, which matches the ligand-receptor pairs in their curated database. The LRPlot() function was used to visualize the differential ligand-receptor pairs as a circos plot from the built-in database of iTALK, which contains a total of 2648 non-redundant and known interacting ligand-receptor pairs classified into 4 categories based on the primary function of the ligand: cytokines, checkpoint proteins, growth factors and others. Red connections indicate gain of interaction, blue or other colors indicate loss of interaction, and the symbolling to provide detailed representation of the changes is described in the key provided in the figure.

#### Transcription Factor-target Gene Regulatory Network Construction

To gain insight into possible regulatory relationships between differentially expressed transcription factors (TF) and their targets, TF-to-gene networks were integrated with DEGs information. Methods were adapted from those previously employed in studies of the effects of shear on cultured endothelial cells(33). Curated and predicted Transcription Factor Targets databases were first downloaded from (https://maayanlab.cloud/Harmonizome/download)(77, 78). The databases include target genes of TF from published ChIP-chip, ChIP-seq, and other TF binding site profiling studies, and TF target genes predicted using known TF binding site motifs. All TF-target databases were then combined to obtain a unique TF-target-gene relationships database. When constructing the DEG-based TF-gene networks, only DEGs that passed certain Fold-Change and p-value thresholds were used. A discrete list of genes of interest can also be used. The networks were generated using R package igraph. The Kamada-Kawai layout algorithm was used to place the vertices on the plane, based on the physical of springs, and the size of the vertices is proportional to the edges it connects. In this layout, the genes with greater number of connections are placed towards the center of the network.

#### Pseudotime Analysis of Endothelial Cells Subclusters

To evaluate possible temporal relationships between the three subclusters of endothelial cells, the Monocle3 R package was used following the published workflow from <u>Monocle 3 (cole-trapnell-lab.github.io)</u>(79). Briefly, the related cell types were re-clustered using the Monocle3 cluster_cells() function, the learn_graph() function was used to generate the principal graph from the reduced dimension space, and then the order_cells() function with root_cells was used to set the EC2 cluster as the start point of the trajectory. The cells were colored with pseudotime inferred and plotted to the UMAP to show the temporal progression of the cells.

#### Differential Blood Flow-related Datasets

Seeking better understanding of blood flow modulation of the endothelial cell transcriptome in the human coronary artery, all endothelial cells were divided into those with low, intermediate or high levels of expression (bottom 1/3, middle 1/3, or top 1/3) of KLF2, which is one of the key TF mediating the anti-atherogenic effects of unidirectional blood flow(42). DEGs were then identified in the KLF2 high versus KLF2 low populations.

To compare the coronary endothelium findings with observations made in an in vivo model of disturbed blood flow, publicly-available single cell RNAseq data for intimal cells in the mouse carotid artery partial ligation model of disturbed blood flow were interrogated(44). The data were downloaded from the NCBI BioProject repository (accession number PRJNA646233). The ligation of major branches of the left carotid artery (LCA) results in disturbed flow and the right carotid artery (RCA) provides the control condition of linear, unidirectional flow(43, 44). The data were preprocessed and scRNA-seq re-analysis was done using the procedures described above for the analysis of the human scRNA-seq data. Cell types were annotated for each cluster using the cell-type information provided(44). For mouse endothelial cells, DEGs were identified between LCA and RCA at either 2 days or 2 weeks following the LCA ligation. To enable the comparison, the human and mouse genes were mapped with homolog gene information using the R package homologene. DEGs from human KLF2 high versus KLF2 low endothelial cells and mouse RCA versus LCA endothelial cells with p< 0.01 were retained and scatterplots with correlations were plotted.

#### Web Portal Development

To facilitate public access and interactive exploration of our study data and results, we developed a user-friendly web portal (https://ai.swmed.edu/Xu-lab-CAD/). The backend infrastructure was implemented using MySQL (https://www.mysql.com) for data management and PHP (https://www.php.net) for dynamic server-side scripting. The database architecture and user interface components were constructed using JavaScript and Bootstrap (https://getbootstrap.com/docs/3.4/javascript/) to ensure responsiveness and cross-platform compatibility. For front-end data visualization, we employed Chart.js and D3.js, enabling dynamic and interactive graphical representations of complex datasets. Additionally, data preprocessing, format conversion, and quality curation were performed using the R programming environment (https://www.r-project.org/), which allowed efficient integration of statistical analyses and visualization outputs into the web interface.

#### Mouse Models

Experiments were performed in male wild-type, Apoe^-/-^, SR-BI^fl/fl^ and HIF-1α^fl/fl^ mice(1, 80), and vascular endothelial cadherin promoter-driven Cre mice (VECad-Cre)(81), or in offspring from their mating. All experiments were approved by the UT Southwestern and the University of Chicago Institutional Animal Care and Use Committees. To evaluate in vivo effects of hypercholesterolemia, beginning at weaning at 4 weeks of age, wild-type mice were fed a standard diet (Teklad Rodent Diet 2018, Intov), and Apoe^-/-^ were fed an atherogenic (D12108C, 20% fat, 1.25% cholesterol, Research Diets Inc.) for 8 weeks. Hypercholesterolemia was also generated by an IV injection of AAV8 encoding a constitutively active form of PCSK9 (5 X 10^11^ genome copies per mouse) at 5 weeks of age and placement on TD96335 (6.2% fat, 1.25% cholesterol, Harlan Laboratories) (1). To increase HIF-1α abundance in vivo, at 8 weeks of age wild-type mice were treated with the HIF prolyl hydroxylase inhibitor dimethyloxalylglycine (DMOG, 100mg/kg IP daily) for 3d and samples were obtained on day 4(37). In the studies of endothelial SR-BI the control mice were Apoe^-/-^;SR-BI^fl/fl^, and the Apoe^-/-^;SR-BI^fl/fl^;VECad-Cre mice deficient selectively in endothelial SR-BI were designated Apoe^-/-^;SR-BI^ΔEC^. HIF-1α^fl/fl^ and VECad-Cre crosses yielded mice lacking HIF-1α in endothelial cells, designated HIF-1α^ΔEC^. In select experiments endothelial cells were isolated from the aorta using rat anti-mouse CD31 (1:500, BD Biosciences cat. No. 553370) and covalently bound sheep anti-rat IgG in magnetic beads (Dynabeads, ThermoFisher)(82).

Studies evaluating the effects of disturbed blood flow on endothelial SR-BI and related mechanisms were performed in the mouse partial carotid artery ligation model(43, 83). Under ketamine and xylazine anesthesia a ventral midline incision was made in the neck, the LCA was exposed, and 3 of the 4 caudal branches of the LCA (left external carotid, internal carotid, and occipital artery) were ligated with 6.0 silk suture, and the superior thyroid artery was left intact. The incision was closed and the mice were allowed to recover. To evaluate changes in endothelial cell gene expression, the control right carotid artery (RCA) and the LCA were harvested 2d post-ligation and intimal RNA was isolated by flushing of the carotid lumen with QIAzol lysis reagent (QIAGEN). Transcript levels were determined and normalized to the geometric mean of three housekeeping genes: β-actin, ubiquitin, and GAPDH. In select studies DiI-labeled native LDL (DiI-LDL) uptake by the aorta was evaluated using our established approach(1). Prior methods for fluorescence imaging of DiI-LDL uptake in the aorta (1) were minimally modified to visualize DiI-LDL in the carotid arteries. Uptake was imaged in the carotid arteries 2d post-ligation in Apoe^-/-^ mice on hypercholesterolemic diet for 1week at 8 weeks of age. DiI-nLDL (100 ul of 1mg/ml) was administered by periorbital injection, and 4h later the carotid arteries were harvested and opened longitudinally to expose the endothelial surface. Following permeabilization with 0.3% Triton X-100 for 30 minutes, tissues were blocked in 2% bovine serum albumin (BSA) for 1 hour at room temperature. Arteries were then incubated overnight at 4 °C with primary antibody against VE-cadherin (Abcam, #ab33168) and Alexa Fluor 488-conjugated anti-rabbit secondary antibody (Abcam, #ab150081) to identify the endothelial cell plasma membranes. After three washes with PBS, tissues were incubated with Alexa Fluor 594-conjugated anti-rabbit secondary antibody (Invitrogen) for 1 hour at room temperature. Subsequently, arteries were counterstained with DAPI (1:1000 dilution) for 10 min and mounted using anti-fade mounting medium (Abcam, UK) with the endothelium facing the coverslip. Fluorescence imaging was performed using a 3i Marianas Spinning Disk Confocal Microscope and collected with SlideBook software (Intelligent Imaging Innovations, CO, USA). DiI detection was quantified in a blinded fashion using Image J.

Studies of atherosclerosis invoked by disturbed flow were performed in the Apoe^-/-^ mouse LCA 1wk post-ligation after hypercholesterolemic diet feeding for 1week at 8 weeks of age. After euthanasia and saline perfusion, aorta, heart, and carotid arteries were isolated en bloc. Heart and carotid arteries were fixed in 4% paraformaldehyde, sequentially immersed in 30% sucrose and optimal cutting temperature compound (OCT):30% sucrose (1:1) mixed solution, and then embedded in OCT. To evaluate atherosclerotic lesion size and visualize lipid accumulation, frozen-embedded samples were sectioned at the mid-portion of the common carotid artery at 8μm thickness and stained with an Oil Red O staining kit (Sigma-Aldrich, MO, USA and ScienCell, CA, USA) according to the manufacturers’ instructions. Lesions were quantified in a blinded fashion using Image J.

#### Quantative RT-PCR

Transcript abundance was evaluated in isolated mouse aortic endothelial cells and cultured human aortic endothelial cells (HAEC) by qRT-PCR using previously established and described methods(84). Primers are listed in Supplementary Table 4.

#### Immunoblotting

Protein abundance was evaluated in harvested mouse aortic endothelial cells and in cultured human aortic endothelial cells (HAEC) by established methods(1). Anti-SR-BI (Abcam, ab52629), anti-actin (Bio-Techne, MAB8929) and anti-GAPDH antibodies (ThermoFisher MA5-15738) were used.

#### Cultured Endothelial Cell Experiments

Experiments were performed in human aortic endothelial cells (HAEC, Lonza) that were maintained in EBM2 medium with 10% (v/v) fetal bovine serum and studied at passages 3–6. SR-BI deletion was accomplished using predesigned and validated shRNA targeting SR-BI packaged in lentivirus (Catalog # <u>VB240723-1488hdy</u>, VectorBuilder). Scrambled shRNA was employed as control intervention (Catalog # VB010000-0009mxc , VectorBuilder). Experiments were performed 48h following lentivirus infection/transduction (MOI 20), and effective knock-down of SR-BI was confirmed by immunoblot analysis. To increase HIF-1α abundance cells were treated with DMOG (1mM) for 24h(85). Changes in HIF-1α protein abundance were determined by ELISA (ab171577, Abcam). In select studies a constitutively-active mutant form of human HIF-1α (Catalog # VB220728-1016rau, VectorBuilder) (86) was introduced into the cells using lentivirus (MOI20), and cell responses were determined 48h later. GFP was introduced by lentivirus (Cat # VB010000-9298rtf, VectorBuilder) to provide a control condition.

#### Endothelial Cell LDL Transcytosis

Transcytosis of LDL was studied in HAEC using our established method(1). Cells were seeded twice, 24 h apart, onto 0.4-μm pore PET Transwell inserts (3610, Corning, Inc.) coated with collagen I (BD Bioscience), with or without lentiviral transduction overnight. Transcytosis was evaluated 48 h later, using fluorescein isothiocyanate (FITC)–dextran (molecular weight, 3,000; Invitrogen) to assess paracellular transport. Cells were incubated simultaneously for 2 h with FITC–dextran and 50 μg/ml DiI–nLDL added to the upper chamber, and the fluorescence intensity of FITC and DiI in the lower chamber was measured using a fluorometer (POLARstar Omega, BMG LABTECH). FITC–dextran transport was 2 to 4%. Infrequently observed inserts displaying paracellular transport more than 5% were discarded. Under control conditions the percent transcytosis of DiI–LDL was approximately 17%, and the data shown are normalized to control values. The validity of the transcytosis assays using DiI–nLDL was previously confirmed using nLDL labelled with iodine-125 by the Bolton–Hunter method(1).

#### ChIP-PCR and ChIP-qPCR

ChIP-PCR and ChIP-qPCR were performed in HAEC to evaluate HIF-1α interaction with the human Scarb1 gene using the High-Sensitivity ChIP Kit (abcam, ab185913). An HIF-1α binding site was identified in intron 1 of Scarb1 in ChIP-seq data from studies of human umbilical vein endothelial cells (HUVEC) placed in normoxic and hypoxic conditions(38). In the present studies, following treatment with DMSO vehicle or DMOG, HAEC were cross-linked for 10min using 1% paraformaldehyde, 1.25M glycine was added to stop the cross-linking reaction, and the cells were harvested and lysed using working lysis buffer containing 6ul/10ml protease inhibitor cocktail (1 x 10^6^ cells in 200ul lysis buffer) to disrupt the cell membrane and release the nuclei. Chromatin was sheared by probe sonication (25% power, 10sec pulses X 4 separated by 30sec on ice). Samples were then centrifuged at 16,000 x g x 3min at 4°C and transfered into strip-wells followed by 1h pre-immunoprecipitation with 100ug/ml Normal Goat IgG Control (R&D AB-108-C) used as negative control or 100ug/ml HIF-1α Ab (R&D AF1935) to capture the antibody-HIF-1α-DNA complexes. Non-specific bound protein-NDA complexes were removed by washing 4 times, and the DNA-HIF-1α complexes were eluted from the beads using DNA release buffer. The eluted DNA was incubated with proteinase K (500ug/ml) at 60°C for 45min to further digest proteins, and the DNA was purified using DNA binding solution, two 90% ethanol washes and column centrifugation. Chromatin from 5X10^5^ cells was used as 10% input. PCR or qPCR was performed to amplify the ChIP products (see Supplementary Table 4 for primer sequences).

#### CRISPR

To test the effect of deleting the HIF-1α binding site in intron 1 of Scarb1, in HAEC the region was excised using CRISPR-Cas9. Two guide RNAs (gRNA) designated gRNA1 and gRNA2 were designed using CRISPOR(87) (see Supplementary Table 2). The dual gRNA and Cas9 were packaged into lentivirus (VectorBuilder VB230802-1416jgv and VB230802-1425dkd) for transduction in HAEC. Deletion of the targeted region was confirmed by PCR (see primer sequences in Supplementary Table 4).

#### Statistical Analysis and Reproducibility

When analyzing the findings in endothelial cells or mice, following normality testing by Shapiro–Wilk test, for normally distributed datasets comparisons between two groups were performed by two-sided Student’s *t* tests, and differences between more than two groups were evaluated by one-way analysis of variance (ANOVA) with Tukey’s post-hoc testing, or by two-way ANOVA with Sidak’s post-hoc testing. In the instances in which normality testing failed, non-parametric analyses were performed between two non-paired groups using Mann–Whitney, between two paired groups using Wilcoxon matched-pairs signed rank test, and between more than two groups by Kruskal–Wallis with Dunn’s post-hoc testing. Findings in cell culture experiments were replicated in two independent experiments. Values shown are mean ± SEM. Significance was accepted at the 0.05 level of probability.

### Data Availability

The raw scRNA-seq data in this study were deposited in GEO under record GSE303311. The secure token for Editors and Reviewers is ebcjysmqppcpdap. There are no restrictions on data availability.

## Acknowledgements

The work was supported by the National Institutes of Health (NIH) grants R01 HL131597, R01 HL144572 and R01 HL114969 (P.W.S.), R01 HL126795 and R01 DK130961 (C.M.), R01 CA245318 and R01 CA258524 (B.L.), R35 HL161244 (Y.F.), R21 CA2733282 (L.X.), the American Heart Association (AHA) grants 23EIA1038679 (Y.F.) and 24POST1198682 (J.Z.), the Sam Day Foundation (L.X.), and the Children’s Cancer Fund (L.X.).

## References

1. Huang L, Chambliss KL, Gao X, Yuhanna IS, Behling-Kelly E, Bergaya S, et al. SR-B1 drives endothelial cell LDL transcytosis via DOCK4 to promote atherosclerosis. Nature. 2019;569(7757):565–9.

2. Moore KJ, Sheedy FJ, and Fisher EA. Macrophages in atherosclerosis: a dynamic balance. Nat Rev Immunol. 2013;13(10):709–21.

3. Pryma CS, Ortega C, Dubland JA, and Francis GA. Pathways of smooth muscle foam cell formation in atherosclerosis. Curr Opin Lipidol. 2019;30(2):117–24.

4. Depuydt MAC, Prange KHM, Slenders L, Ord T, Elbersen D, Boltjes A, et al. Microanatomy of the Human Atherosclerotic Plaque by Single-Cell Transcriptomics. Circ Res. 2020;127(11):1437–55.

5. Turner AW, Hu SS, Mosquera JV, Ma WF, Hodonsky CJ, Wong D, et al. Single-nucleus chromatin accessibility profiling highlights regulatory mechanisms of coronary artery disease risk. Nat Genet. 2022;54(6):804–16.

6. Subramanian Vignesh K, and Deepe GS, Jr. Metallothioneins: Emerging Modulators in Immunity and Infection. Int J Mol Sci. 2017;18(10).

7. Kwan WH, van der Touw W, and Heeger PS. Complement regulation of T cell immunity. Immunol Res. 2012;54(1-3):247–53.

8. Kanbar JN, Ma S, Kim ES, Kurd NS, Tsai MS, Tysl T, et al. The long noncoding RNA Malat1 regulates CD8+ T cell differentiation by mediating epigenetic repression. J Exp Med. 2022;219(6).

9. Selathurai A, Deswaerte V, Kanellakis P, Tipping P, Toh BH, Bobik A, et al. Natural killer (NK) cells augment atherosclerosis by cytotoxic-dependent mechanisms. Cardiovasc Res. 2014;102(1):128–37.

10. Zaiss DMW, Gause WC, Osborne LC, and Artis D. Emerging functions of amphiregulin in orchestrating immunity, inflammation, and tissue repair. Immunity. 2015;42(2):216–26.

11. Sage AP, Tsiantoulas D, Binder CJ, and Mallat Z. The role of B cells in atherosclerosis. Nat Rev Cardiol. 2019;16(3):180–96.

12. Chen R, McVey DG, Shen D, Huang X, and Ye S. Phenotypic Switching of Vascular Smooth Muscle Cells in Atherosclerosis. J Am Heart Assoc. 2023;12(20):e031121.

13. Rasbach E, Splitthoff P, Bonaterra GA, Schwarz A, Mey L, Schwarzbach H, et al. PACAP deficiency aggravates atherosclerosis in ApoE deficient mice. Immunobiology. 2019;224(1):124–32.

14. Nelson CP, Goel A, Butterworth AS, Kanoni S, Webb TR, Marouli E, et al. Association analyses based on false discovery rate implicate new loci for coronary artery disease. Nat Genet. 2017;49(9):1385–91.

15. Jiang XC, and Yu Y. The Role of Phospholipid Transfer Protein in the Development of Atherosclerosis. Curr Atheroscler Rep. 2021;23(3):9.

16. O’Brien KD, Vuletic S, McDonald TO, Wolfbauer G, Lewis K, Tu AY, et al. Cell-associated and extracellular phospholipid transfer protein in human coronary atherosclerosis. Circulation. 2003;108(3):270–4.

17. Zhang F, Fu X, Kataoka M, Liu N, Wang Y, Gao F, et al. Long noncoding RNA Cfast regulates cardiac fibrosis. Mol Ther Nucleic Acids. 2021;23:377–92.

18. Li L, Zhang Q, Lei X, Huang Y, and Hu J. MAP4 as a New Candidate in Cardiovascular Disease. Front Physiol. 2020;11:1044.

19. Dennis J, Johnson CY, Adediran AS, de Andrade M, Heit JA, Morange PE, et al. The endothelial protein C receptor (PROCR) Ser219Gly variant and risk of common thrombotic disorders: a HuGE review and meta-analysis of evidence from observational studies. Blood. 2012;119(10):2392–400.

20. Li R, Paul A, Ko KW, Sheldon M, Rich BE, Terashima T, et al. Interleukin-7 induces recruitment of monocytes/macrophages to endothelium. Eur Heart J. 2012;33(24):3114–23.

21. Cardilo-Reis L, Gruber S, Schreier SM, Drechsler M, Papac-Milicevic N, Weber C, et al. Interleukin-13 protects from atherosclerosis and modulates plaque composition by skewing the macrophage phenotype. EMBO Mol Med. 2012;4(10):1072–86.

22. Edsfeldt A, Singh P, Matthes F, Tengryd C, Cavalera M, Bengtsson E, et al. Transforming growth factor-beta2 is associated with atherosclerotic plaque stability and lower risk for cardiovascular events. Cardiovasc Res. 2023;119(11):2061–73.

23. Murakami M, Nguyen LT, Zhuang ZW, Moodie KL, Carmeliet P, Stan RV, et al. The FGF system has a key role in regulating vascular integrity. J Clin Invest. 2008;118(10):3355–66.

24. Vuong JT, Stein-Merlob AF, Nayeri A, Sallam T, Neilan TG, and Yang EH. Immune Checkpoint Therapies and Atherosclerosis: Mechanisms and Clinical Implications: JACC State-of-the-Art Review. J Am Coll Cardiol. 2022;79(6):577–93.

25. Wang J, Zhou X, Su Y, Chai D, Ruan Y, and Wang J. Association between haptoglobin polymorphism and coronary artery disease: a meta-analysis. Front Genet. 2024;15:1434975.

26. Ivetic A, Hoskins Green HL, and Hart SJ. L-selectin: A Major Regulator of Leukocyte Adhesion, Migration and Signaling. Front Immunol. 2019;10:1068.

27. Cheent KS, Jamil KM, Cassidy S, Liu M, Mbiribindi B, Mulder A, et al. Synergistic inhibition of natural killer cells by the nonsignaling molecule CD94. Proc Natl Acad Sci U S A. 2013;110(42):16981–6.

28. Yao Y, Bennett BJ, Wang X, Rosenfeld ME, Giachelli C, Lusis AJ, et al. Inhibition of bone morphogenetic proteins protects against atherosclerosis and vascular calcification. Circ Res. 2010;107(4):485–94.

29. Gu N, Dong Y, Tian Y, Di Z, Liu Z, Chang M, et al. Anti-apoptotic and angiogenic effects of intelectin-1 in rat cerebral ischemia. Brain Res Bull. 2017;130:27–35.

30. Wei M, Pan H, and Guo K. Association Between Plasma ADAMTS-9 Levels and Severity of Coronary Artery Disease. Angiology. 2021;72(4):371–80.

31. Koo BH, Coe DM, Dixon LJ, Somerville RP, Nelson CM, Wang LW, et al. ADAMTS9 is a cell-autonomously acting, anti-angiogenic metalloprotease expressed by microvascular endothelial cells. Am J Pathol. 2010;176(3):1494–504.

32. Wan W, Liu Q, Lionakis MS, Marino AP, Anderson SA, Swamydas M, et al. Atypical chemokine receptor 1 deficiency reduces atherogenesis in ApoE-knockout mice. Cardiovasc Res. 2015;106(3):478–87.

33. Maurya MR, Gupta S, Li JY, Ajami NE, Chen ZB, Shyy JY, et al. Longitudinal shear stress response in human endothelial cells to atheroprone and atheroprotective conditions. Proc Natl Acad Sci U S A. 2021;118(4).

34. Abdelzaher AF, Al-Musawi AF, Ghosh P, Mayo ML, and Perkins EJ. Transcriptional Network Growing Models Using Motif-Based Preferential Attachment. Front Bioeng Biotechnol. 2015;3:157.

35. Thomas C, Leleu D, and Masson D. Cholesterol and HIF-1alpha: Dangerous Liaisons in Atherosclerosis. Front Immunol. 2022;13:868958.

36. Koyasu S, Kobayashi M, Goto Y, Hiraoka M, and Harada H. Regulatory mechanisms of hypoxia-inducible factor 1 activity: Two decades of knowledge. Cancer Sci. 2018;109(3):560–71.

37. Baader E, Tschank G, Baringhaus KH, Burghard H, and Gunzler V. Inhibition of prolyl 4-hydroxylase by oxalyl amino acid derivatives in vitro, in isolated microsomes and in embryonic chicken tissues. Biochem J. 1994;300 ( Pt 2)(Pt 2):525–30.

38. Mimura I, Nangaku M, Kanki Y, Tsutsumi S, Inoue T, Kohro T, et al. Dynamic change of chromatin conformation in response to hypoxia enhances the expression of GLUT3 (SLC2A3) by cooperative interaction of hypoxia-inducible factor 1 and KDM3A. Mol Cell Biol. 2012;32(15):3018–32.

39. Grandoch M, Kohlmorgen C, Melchior-Becker A, Feldmann K, Homann S, Muller J, et al. Loss of Biglycan Enhances Thrombin Generation in Apolipoprotein E-Deficient Mice: Implications for Inflammation and Atherosclerosis. Arterioscler Thromb Vasc Biol. 2016;36(5):e41–50.

40. Soubeyrand S, Lau P, Nikpay M, Dang AT, and McPherson R. Common Polymorphism That Protects From Cardiovascular Disease Increases Fibronectin Processing and Secretion. Circ Genom Precis Med. 2022;15(2):e003428.

41. Mahmoodi M, Mirzarazi Dahagi E, Nabavi MH, Penalva YCM, Gosaine A, Murshed M, et al. Circulating plasma fibronectin affects tissue insulin sensitivity, adipocyte differentiation, and transcriptional landscape of adipose tissue in mice. Physiol Rep. 2024;12(14):e16152.

42. Tamargo IA, Baek KI, Kim Y, Park C, and Jo H. Flow-induced reprogramming of endothelial cells in atherosclerosis. Nat Rev Cardiol. 2023;20(11):738–53.

43. Nam D, Ni CW, Rezvan A, Suo J, Budzyn K, Llanos A, et al. Partial carotid ligation is a model of acutely induced disturbed flow, leading to rapid endothelial dysfunction and atherosclerosis. Am J Physiol Heart Circ Physiol. 2009;297(4):H1535–43.

44. Andueza A, Kumar S, Kim J, Kang DW, Mumme HL, Perez JI, et al. Endothelial Reprogramming by Disturbed Flow Revealed by Single-Cell RNA and Chromatin Accessibility Study. Cell Rep. 2020;33(11):108491.

45. Zeng H, Qin L, Zhao D, Tan X, Manseau EJ, Van Hoang M, et al. Orphan nuclear receptor TR3/Nur77 regulates VEGF-A-induced angiogenesis through its transcriptional activity. J Exp Med. 2006;203(3):719–29.

46. Furst R, Schroeder T, Eilken HM, Bubik MF, Kiemer AK, Zahler S, et al. MAPK phosphatase-1 represents a novel anti-inflammatory target of glucocorticoids in the human endothelium. FASEB J. 2007;21(1):74–80.

47. Maleszewska M, Gjaltema RA, Krenning G, and Harmsen MC. Enhancer of zeste homolog-2 (EZH2) methyltransferase regulates transgelin/smooth muscle-22alpha expression in endothelial cells in response to interleukin-1beta and transforming growth factor-beta2. Cell Signal. 2015;27(8):1589–96.

48. Ke K, Wu Z, Lin J, Lin L, Huang N, and Yang W. Increased Expression of CD74 in Atherosclerosis Associated with Inflammatory Responses of Endothelial Cells and Macrophages. Biochem Genet. 2024;62(1):294–310.

49. Estevao C, Bowers CE, Luo D, Sarker M, Hoeh AE, Frudd K, et al. CCL4 induces inflammatory signalling and barrier disruption in the neurovascular endothelium. Brain Behav Immun Health. 2021;18:100370.

50. Schulkens IA, Castricum KC, Weijers EM, Koolwijk P, Griffioen AW, and Thijssen VL. Expression, regulation and function of human metallothioneins in endothelial cells. J Vasc Res. 2014;51(3):231–8.

51. Li Z, Qin Z, Kong X, Chen B, Hu W, Lin Z, et al. CCL14 exacerbates intraplaque vulnerability by promoting neovascularization in the human carotid plaque. J Stroke Cerebrovasc Dis. 2022;31(10):106670.

52. Schake MA, McCue IS, Curtis ET, Ripperda TJ, Jr., Harvey S, Hackfort BT, et al. Restoration of normal blood flow in atherosclerotic arteries promotes plaque stabilization. *iScience.* 2023;26(6):106760.

53. Cait J, Hughes MR, Zeglinski MR, Chan AW, Osterhof S, Scott RW, et al. Podocalyxin is required for maintaining blood-brain barrier function during acute inflammation. Proc Natl Acad Sci U S A. 2019;116(10):4518–27.

54. Porras G, Ayuso MS, and Gonzalez-Manchon C. Leukocyte-endothelial cell interaction is enhanced in podocalyxin-deficient mice. Int J Biochem Cell Biol. 2018;99:72–9.

55. Kielczewski JL, Jarajapu YP, McFarland EL, Cai J, Afzal A, Li Calzi S, et al. Insulin-like growth factor binding protein-3 mediates vascular repair by enhancing nitric oxide generation. Circ Res. 2009;105(9):897–905.

56. Kujovich JL. Factor V Leiden thrombophilia. Genet Med. 2011;13(1):1–16.

57. Schwarz N, Muller J, McRae HL, Reda S, Pezeshkpoor B, Oldenburg J, et al. Endothelium Modulates the Prothrombotic Phenotype of Factor V Leiden: Evidence From an Ex Vivo Model. Arterioscler Thromb Vasc Biol. 2025;45(3):412–23.

58. Hazell GG, Peachey AM, Teasdale JE, Sala-Newby GB, Angelini GD, Newby AC, et al. PI16 is a shear stress and inflammation-regulated inhibitor of MMP2. Sci Rep. 2016;6:39553.

59. Schwenke DC, and Carew TE. Quantification in vivo of increased LDL content and rate of LDL degradation in normal rabbit aorta occurring at sites susceptible to early atherosclerotic lesions. Circ Res. 1988;62(4):699–710.

60. Chatzizisis YS, Coskun AU, Jonas M, Edelman ER, Feldman CL, and Stone PH. Role of endothelial shear stress in the natural history of coronary atherosclerosis and vascular remodeling: molecular, cellular, and vascular behavior. J Am Coll Cardiol. 2007;49(25):2379–93.

61. Alloza I, Goikuria H, Idro JL, Trivino JC, Fernandez Velasco JM, Elizagaray E, et al. RNAseq based transcriptomics study of SMCs from carotid atherosclerotic plaque: BMP2 and IDs proteins are crucial regulators of plaque stability. Sci Rep. 2017;7(1):3470.

62. Wirka RC, Wagh D, Paik DT, Pjanic M, Nguyen T, Miller CL, et al. Atheroprotective roles of smooth muscle cell phenotypic modulation and the TCF21 disease gene as revealed by single-cell analysis. Nat Med. 2019;25(8):1280–9.

63. Fernandez DM, Rahman AH, Fernandez NF, Chudnovskiy A, Amir ED, Amadori L, et al. Single-cell immune landscape of human atherosclerotic plaques. Nat Med. 2019;25(10):1576–88.

64. Hu Z, Liu W, Hua X, Chen X, Chang Y, Hu Y, et al. Single-Cell Transcriptomic Atlas of Different Human Cardiac Arteries Identifies Cell Types Associated With Vascular Physiology. Arterioscler Thromb Vasc Biol. 2021;41(4):1408–27.

65. Kalluri AS, Vellarikkal SK, Edelman ER, Nguyen L, Subramanian A, Ellinor PT, et al. Single-Cell Analysis of the Normal Mouse Aorta Reveals Functionally Distinct Endothelial Cell Populations. Circulation. 2019;140(2):147–63.

66. Mao A, Zhang K, Kan H, Gao M, Wang Z, Zhou T, et al. Single-Cell RNA-Seq Reveals Coronary Heterogeneity and Identifies CD133(+)TRPV4(high) Endothelial Subpopulation in Regulating Flow-Induced Vascular Tone in Mice. Arterioscler Thromb Vasc Biol. 2024;44(3):653–65.

67. Foteinos G, Hu Y, Xiao Q, Metzler B, and Xu Q. Rapid endothelial turnover in atherosclerosis-prone areas coincides with stem cell repair in apolipoprotein E-deficient mice. Circulation. 2008;117(14):1856–63.

68. Davies PF, Civelek M, Fang Y, and Fleming I. The atherosusceptible endothelium: endothelial phenotypes in complex haemodynamic shear stress regions in vivo. Cardiovasc Res. 2013;99(2):315–27.

69. Pan B, Ma Y, Ren H, He Y, Wang Y, Lv X, et al. Diabetic HDL is dysfunctional in stimulating endothelial cell migration and proliferation due to down regulation of SR-BI expression. PLoS One. 2012;7(11):e48530.

70. Ghaffari S, Naderi Nabi F, Sugiyama MG, and Lee WL. Estrogen Inhibits LDL (Low-Density Lipoprotein) Transcytosis by Human Coronary Artery Endothelial Cells via GPER (G-Protein-Coupled Estrogen Receptor) and SR-BI (Scavenger Receptor Class B Type 1). Arterioscler Thromb Vasc Biol. 2018;38(10):2283–94.

71. Boren J, Chapman MJ, Krauss RM, Packard CJ, Bentzon JF, Binder CJ, et al. Low-density lipoproteins cause atherosclerotic cardiovascular disease: pathophysiological, genetic, and therapeutic insights: a consensus statement from the European Atherosclerosis Society Consensus Panel. Eur Heart J. 2020;41(24):2313–30.

72. van Veelen A, van der Sangen NMR, Henriques JPS, and Claessen B. Identification and treatment of the vulnerable coronary plaque. Rev Cardiovasc Med. 2022;23(1):39.

73. Virmani R, Kolodgie FD, Burke AP, Farb A, and Schwartz SM. Lessons from sudden coronary death: a comprehensive morphological classification scheme for atherosclerotic lesions. Arterioscler Thromb Vasc Biol. 2000;20(5):1262–75.

74. Stary HC, Chandler AB, Dinsmore RE, Fuster V, Glagov S, Insull W, Jr., et al. A definition of advanced types of atherosclerotic lesions and a histological classification of atherosclerosis. A report from the Committee on Vascular Lesions of the Council on Arteriosclerosis, American Heart Association. Circulation. 1995;92(5):1355–74.

75. Stuart T, Butler A, Hoffman P, Hafemeister C, Papalexi E, Mauck WM, 3rd, et al. Comprehensive Integration of Single-Cell Data. Cell. 2019;177(7):1888–902 e21.

76. Wang Y, Wang R, Zhang S, Song S, Jiang C, Han G, et al. iTALK: an R Package to Characterize and Illustrate Intercellular Communication. *bioRxiv*. 2019:507871.

77. Diamant I, Clarke DJB, Evangelista JE, Lingam N, and Ma’ayan A. Harmonizome 3.0: integrated knowledge about genes and proteins from diverse multi-omics resources. Nucleic Acids Res. 2025;53(D1):D1016–D28.

78. Rouillard AD, Gundersen GW, Fernandez NF, Wang Z, Monteiro CD, McDermott MG, et al. The harmonizome: a collection of processed datasets gathered to serve and mine knowledge about genes and proteins. Database (Oxford). 2016;2016.

79. Trapnell C, Cacchiarelli D, Grimsby J, Pokharel P, Li S, Morse M, et al. The dynamics and regulators of cell fate decisions are revealed by pseudotemporal ordering of single cells. Nat Biotechnol. 2014;32(4):381–6.

80. Ryan HE, Poloni M, McNulty W, Elson D, Gassmann M, Arbeit JM, et al. Hypoxia-inducible factor-1alpha is a positive factor in solid tumor growth. Cancer Res. 2000;60(15):4010–5.

81. Alva JA, Zovein AC, Monvoisin A, Murphy T, Salazar A, Harvey NL, et al. VE-Cadherin-Cre-recombinase transgenic mouse: a tool for lineage analysis and gene deletion in endothelial cells. Dev Dyn. 2006;235(3):759–67.

82. Sacharidou A, Chambliss K, Peng J, Barrera J, Tanigaki K, Luby-Phelps K, et al. Endothelial ERalpha promotes glucose tolerance by enhancing endothelial insulin transport to skeletal muscle. Nat Commun. 2023;14(1):4989.

83. Yeh CF, Cheng SH, Lin YS, Shentu TP, Huang RT, Zhu J, et al. Targeting mechanosensitive endothelial TXNDC5 to stabilize eNOS and reduce atherosclerosis in vivo. Sci Adv. 2022;8(3):eabl8096.

84. Umetani M, Ghosh P, Ishikawa T, Umetani J, Ahmed M, Mineo C, et al. The cholesterol metabolite 27-hydroxycholesterol promotes atherosclerosis via proinflammatory processes mediated by estrogen receptor alpha. Cell Metab. 2014;20(1):172–82.

85. Asikainen TM, Schneider BK, Waleh NS, Clyman RI, Ho WB, Flippin LA, et al. Activation of hypoxia-inducible factors in hyperoxia through prolyl 4-hydroxylase blockade in cells and explants of primate lung. Proc Natl Acad Sci U S A. 2005;102(29):10212–7.

86. Kelly BD, Hackett SF, Hirota K, Oshima Y, Cai Z, Berg-Dixon S, et al. Cell type-specific regulation of angiogenic growth factor gene expression and induction of angiogenesis in nonischemic tissue by a constitutively active form of hypoxia-inducible factor 1. Circ Res. 2003;93(11):1074–81.

87. Concordet JP, and Haeussler M. CRISPOR: intuitive guide selection for CRISPR/Cas9 genome editing experiments and screens. Nucleic Acids Res. 2018;46(W1):W242–W5.

