## Extended Data for "Coronary Artery Disease Transcriptomics Reveals Two Drivers of the Endothelial Cell SR-BI Expression and LDL Transport that Underlie Atherosclerosis"

Extended Data Figure 1

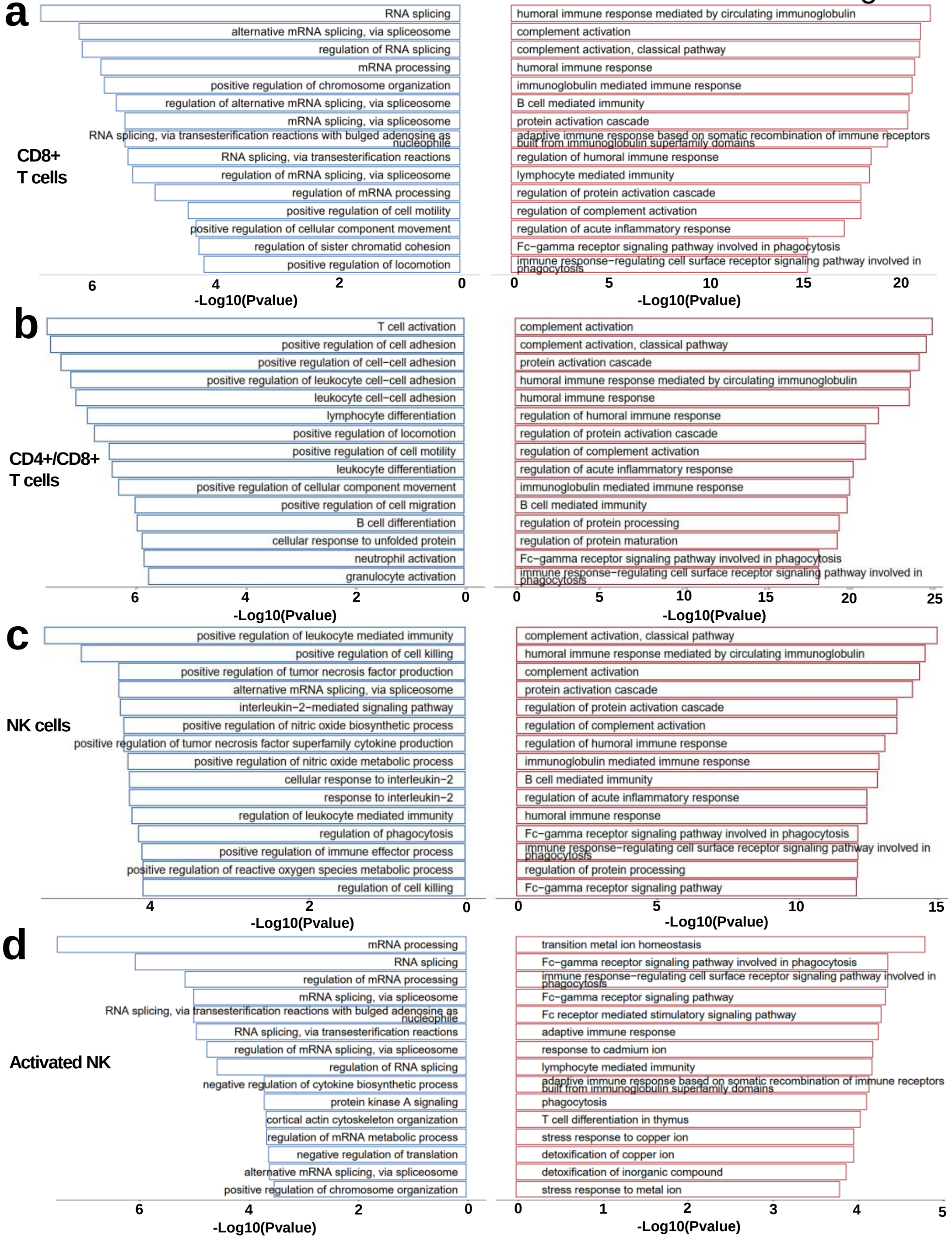


**Extended Data Fig. 1: Functional pathways of genes in CD8+ T cells, CD4+/CD8+ T cells, NK cells and activated NK cells differ in coronary artery segments lacking versus bearing atheroma. a-d**, Top 15 pathways that are more prevalent (red) or less prevalent (blue) in atheroma-containing segments compared to atheroma-free segments in CD8+ T cells (**a**), CD4+/CD8+ T cells (**b**), NK cells (**c**) and activated NK cells (**d**).

Extended Data Figure 2

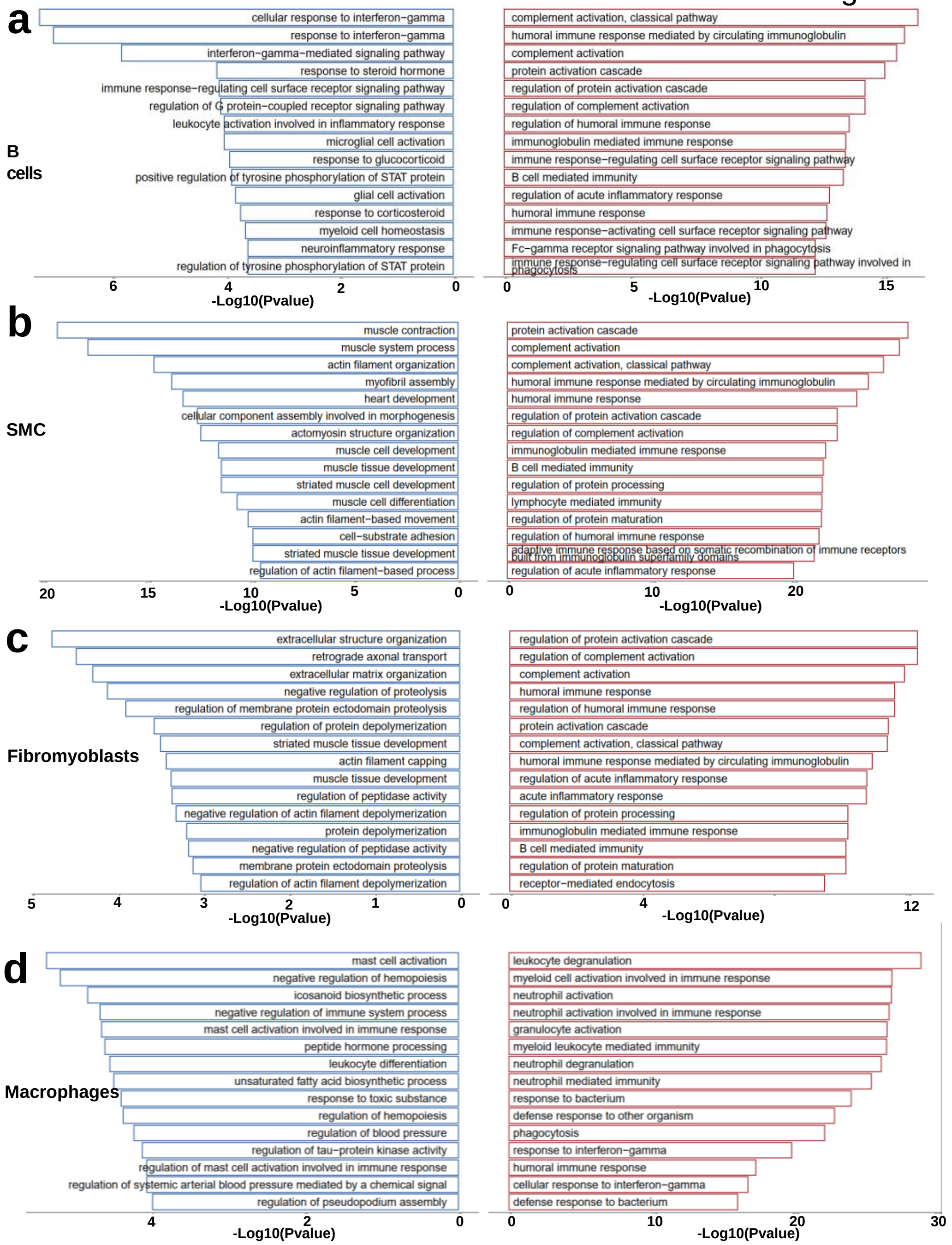


**Extended Data Fig. 2: Functional pathways of genes in B cells, vascular smooth muscle cells, fibromyoblasts and macrophages differ in coronary artery segments lacking versus bearing atheroma. a-d**, Top 15 pathways that are more prevalent (red) or less prevalent (blue) in atheroma-containing segments compared to atheroma-free segments in B cells (**a**), vascular smooth muscle cells (SMC, **b**), fibromyoblasts (**c**) and macrophages (**d**).

Extended Data
Figure 3

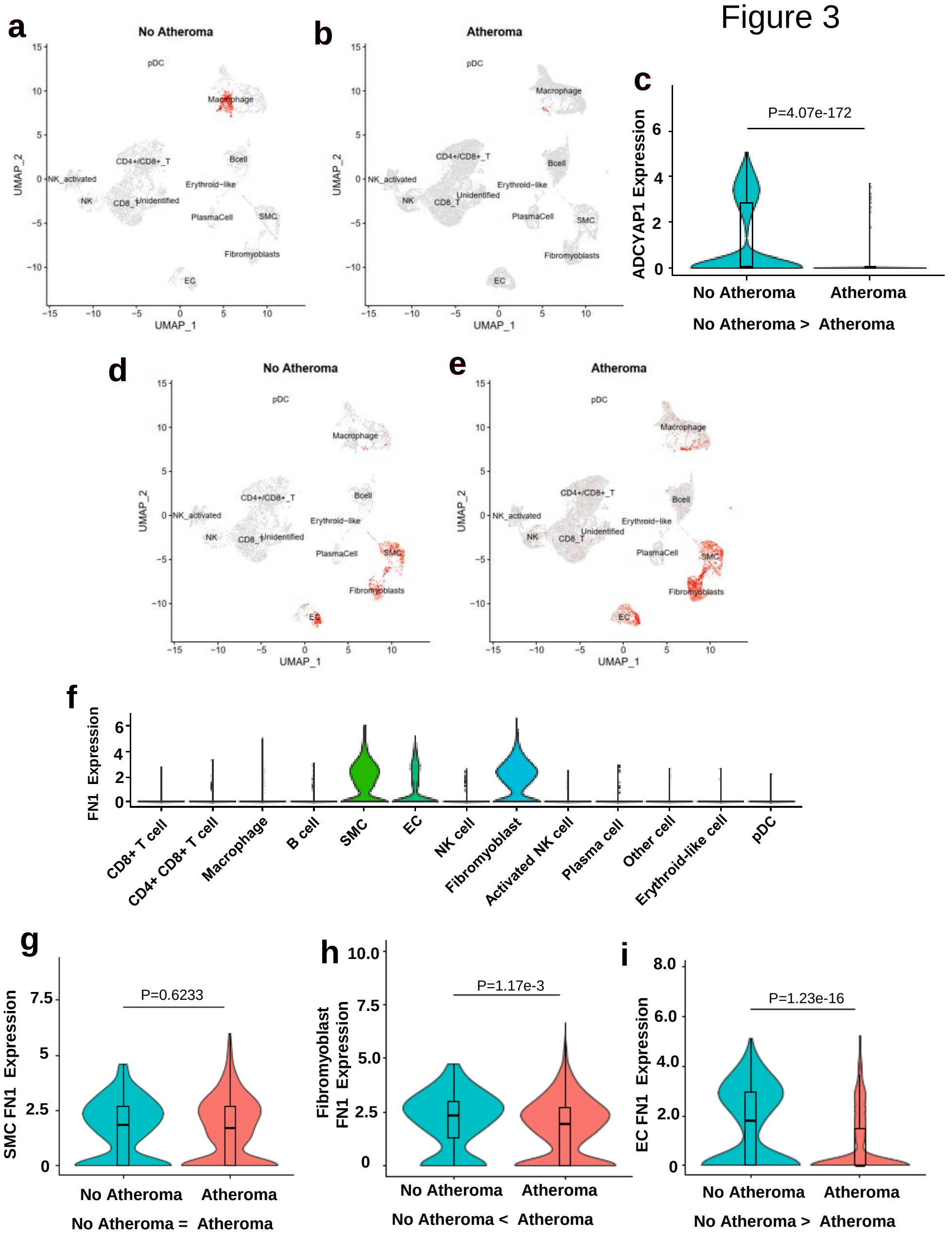


**Extended Data Fig. 3: Web utility findings for ADCYAP1 and FN1 in coronary artery segments lacking versus bearing atheroma. a-c**, ADCYAP1 expression. UMAP plots revealing relative cell type-specific expression of ADCYAP1 in non-atheroma (**a**) and atheroma-containing segments (**b**), and violin plots of ADCYAP1 expression in macrophages in non-atheroma versus atheroma segments (**c**). **d-i**, FN1 expression. UMAP plots revealing relative cell type-specific expression of FN1 in non-atheroma (**d**) and atheroma-containing segments (**e**), violin plots of FN1 expression in all 13 vascular cell types (**f**), and violin plots of FN1 expression in non-atheroma versus atheroma segments in SMC (**g**), fibromyoblasts (**h**) and endothelial cells (EC, **i**).

### Extended Data Figure 4

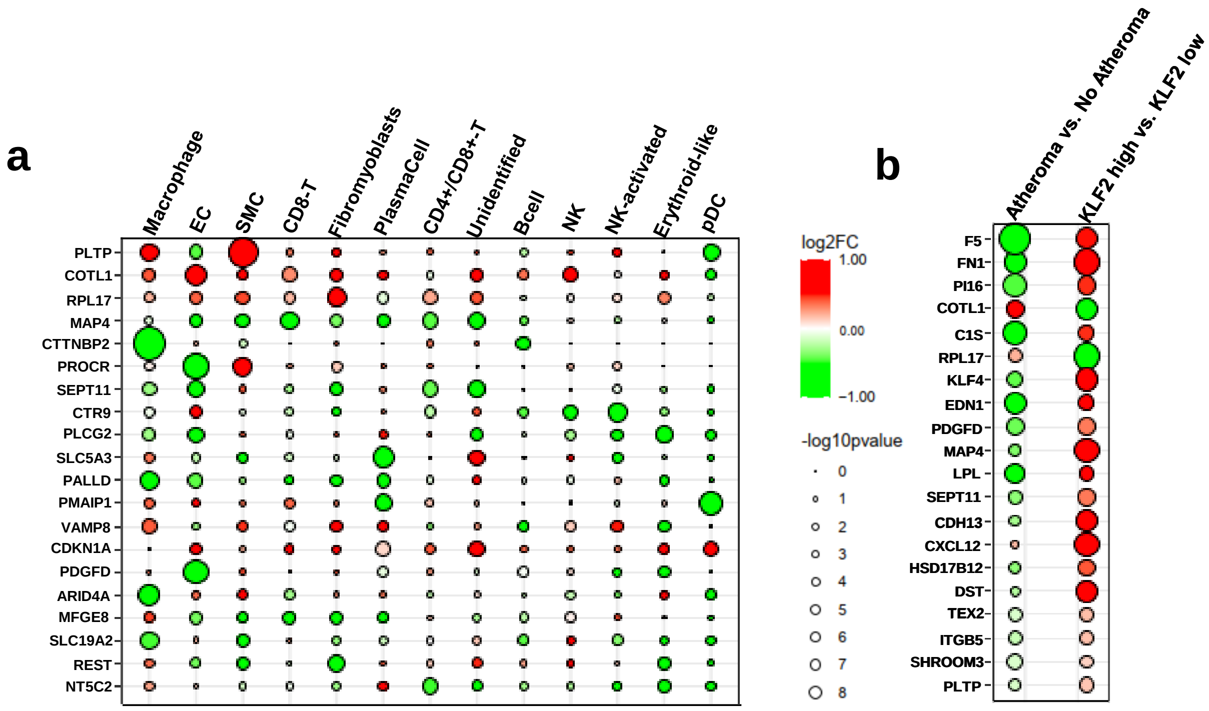


**Extended Data Fig. 4: Relative expression of GWAS-identified CAD risk genes. a,** Dot plot of CAD risk gene expression in 13 specific cell types in non-atheroma versus atheroma coronary artery segments. Decreased expression in atheroma is indicated in green, and increased expression in atheroma is indicated in red. The genes are ranked based on the sum of the negative log 10 p values across all cell types, and the top 20 genes are shown. **b**, Dot plot of CAD risk gene expression in endothelial cells, in atheroma versus non-atheroma-associated endothelial cells (left), and in KLF2 high versus KLF2 low endothelial cell populations (right). The genes are ranked based on the product of the negative log 10 p values in the two comparisons, and the top 20 genes are shown.

Extended Data Figure 5

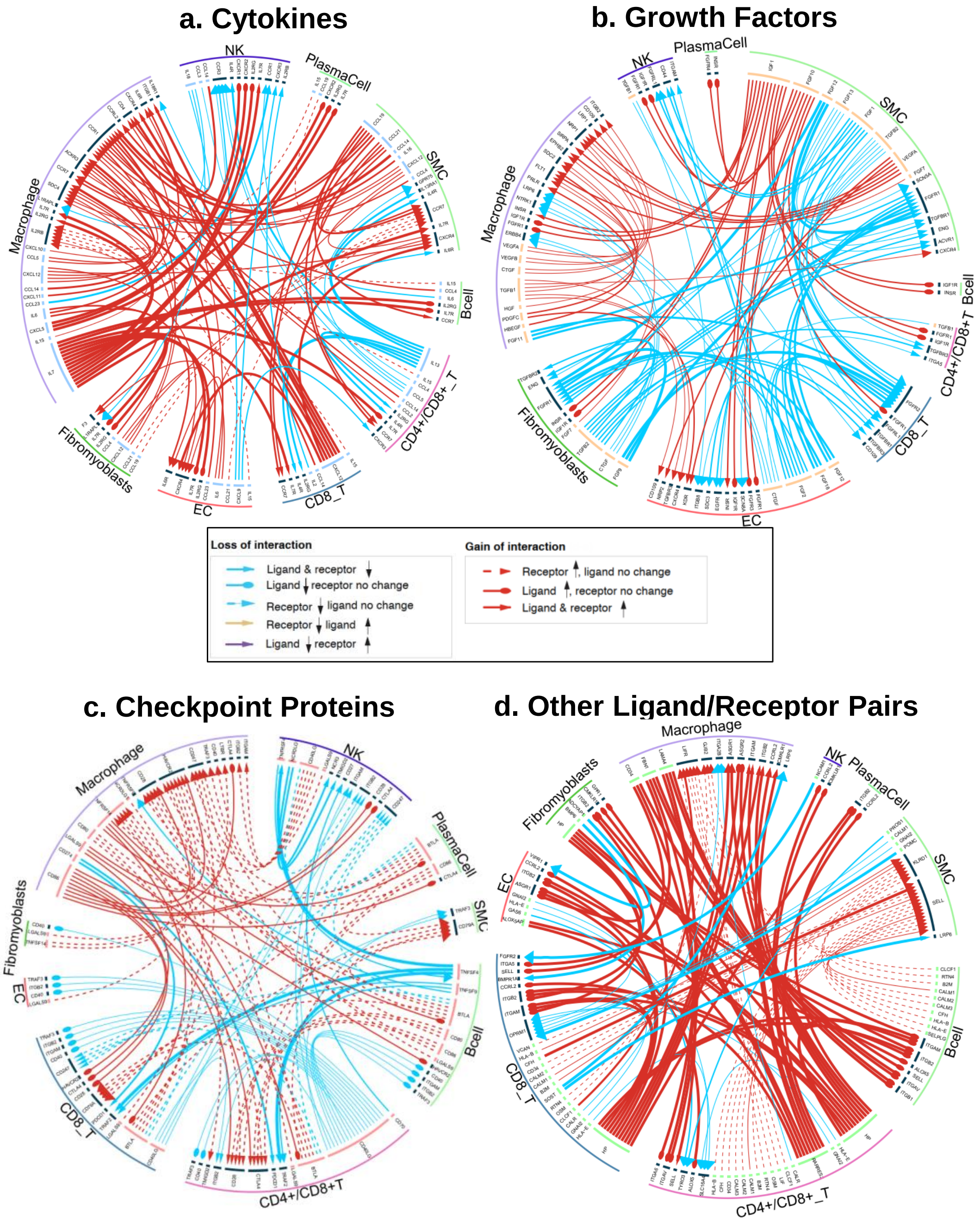


**Extended Data Fig. 5: Cell-to-cell communication changes during the progression of atherosclerosis.** Circos plots are shown for communication related to cytokines (**a**), growth factors (**b**), checkpoint proteins (**c**) and other ligand and receptor autocrine or paracrine interactions (**d**). The circos plots were generated using DEGs to evaluate possible gains of interaction (coded in red), with ligand and/or receptor upregulated in the atheroma segment cells, and possible losses of interaction (coded in blue), with ligand and/or receptor downregulated in atheroma-associated cells.

Extended Data Figure 6

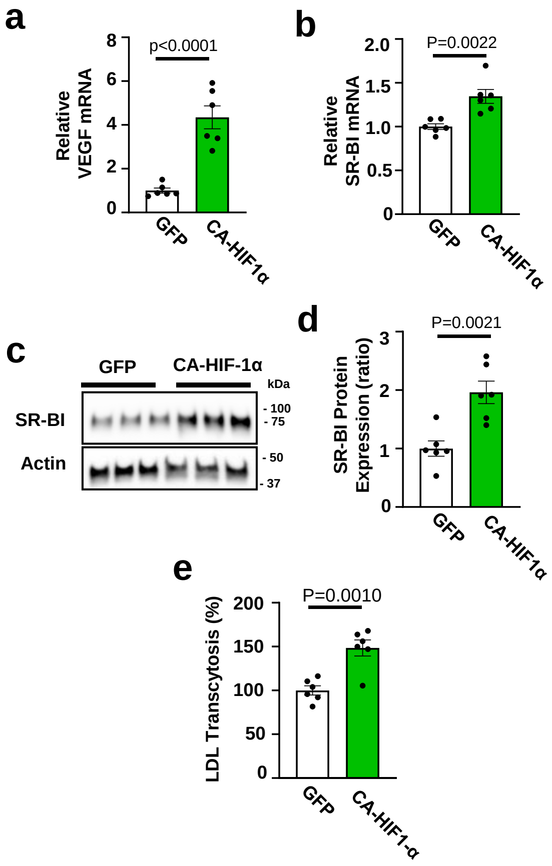


**Extended Data Fig. 6: Endothelial cell SR-BI is upregulated by constitutively-active HIF-1α.** A constitutively-active mutant form of human HIF-1α (CA-HIF-1α) or GFP control was introduced into HAEC using lentivirus, and cell responses were determined 48h later. **a**, Expression of the known HIF-1α target gene VEGF was upregulated. **b-d**, CA-HIF-1α caused an increase in SR-BI mRNA (**b**) and protein abundance (**c,d**). Representative immunoblot showing 3 samples per group is in **c**, and summary data are in **d**. **e**, CA-HIF-1α also caused an increase in the transcytosis of DiI-labeled LDL. Data are mean±SEM. In **a,b,d** and **e,** p values by two-tailed unpaired Student's t test are shown.

Extended Data Figure 7

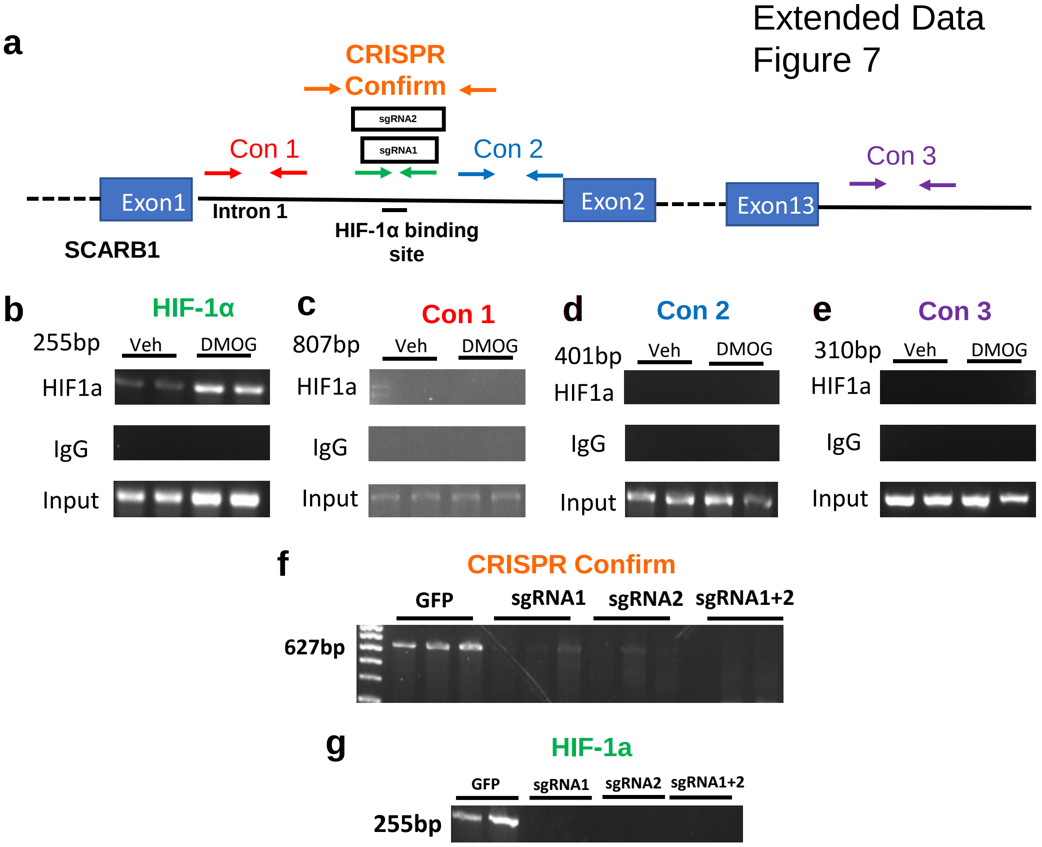


**Extended Data Fig. 7: CRISPR-Cas9 deletion of the HIF-1α binding site in Scarb1 intron 1.** **a**. Schematic of HIF-1α binding site in intron 1 of human Scarb1, location of two sgRNAs, and PCR primers for evaluating the HIF-1α binding site, two control regions in intron 1 (Con1 and Con2), a third remote control region (Con3), and the CRISPR-Cas9 deletion. **b-e**, ChIP-PCR was performed evaluating HIF-1α binding to the predicted HIF-1a binding site (**b**) and the three control sites (**c-e**) 24h following HAEC treatment with DMSO vehicle control or DMOG. The ChIP-PCR was performed with both anti-HIF-1α antibody and an unrelated IgG control. **f**, Evaluation of the CRISPR-Cas9 deletion of the targeted region following introduction of sgRNA1, sgRNA2, or both sgRNA1 and sgRNA2. **g**, Parallel evaluation of deletion of the predicted HIF-1α binding site.

### Extended Data Figure 8

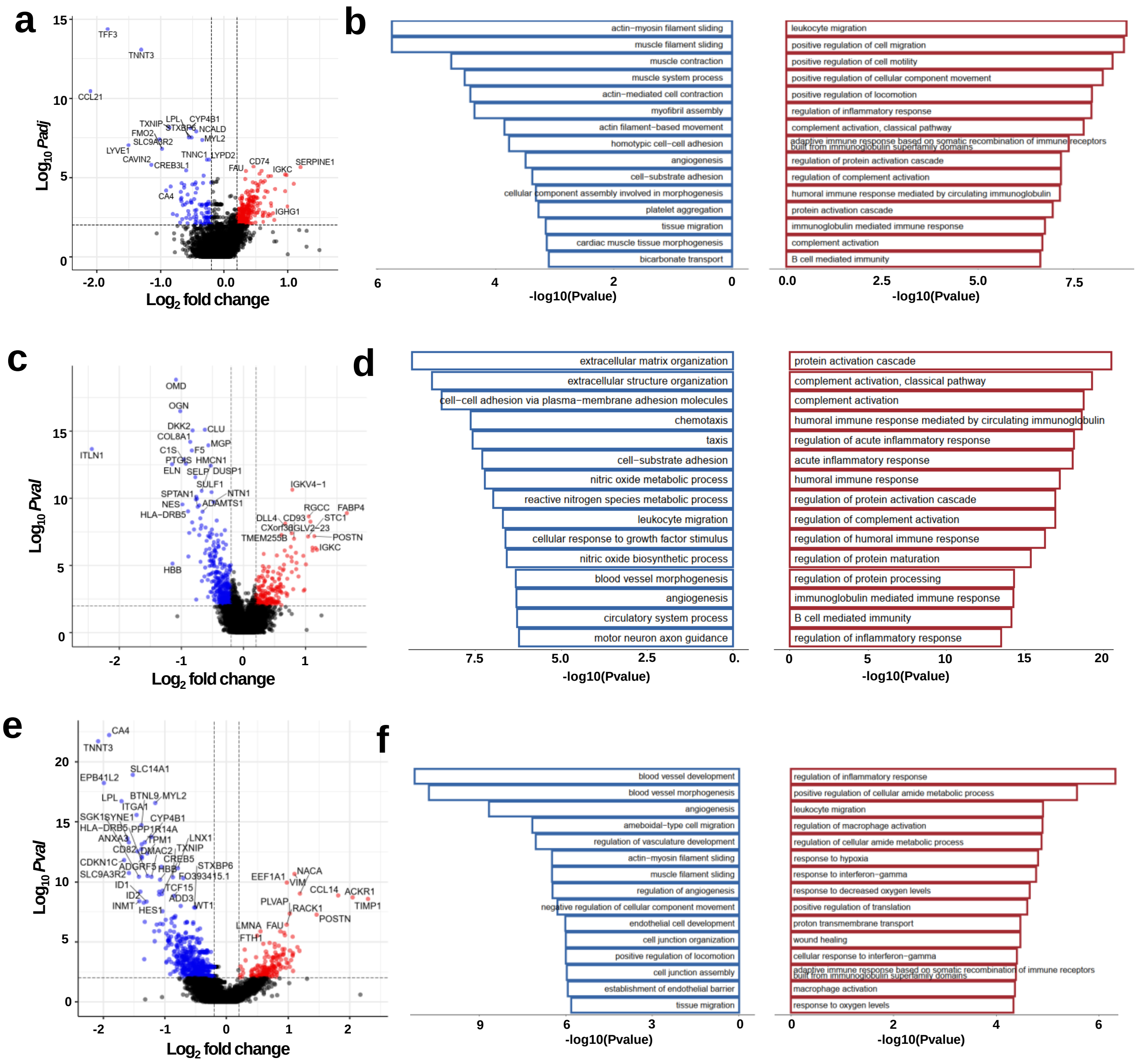

**Extended Data Fig. 8: Transcripts change in luminal endothelial cell subtypes with atheroma formation.** **a,** Volcano plot depicting differentially-expressed genes (DEGs) in EC0 endothelium in atheroma-containing segments compared to atheroma-free segments. Log 2 fold-change >0.2, and adjusted p<0.01. Red dots indicate genes upregulated in EC0 in atheroma bearing segments and blue dots indicate genes downregulated in EC0 associated with atheroma. **b**, Top 15 pathways that are more prevalent (red) or less prevalent (blue) in EC0 in atheroma-containing segments compared to atheroma-free coronary artery. **c**, Volcano plot depicting DEGs in EC1 endothelium in atheroma-containing segments compared to atheroma-free segments. **d**, Top 15 pathways that are more prevalent or less prevalent in EC1 in atheroma-containing compared to atheroma-free segments. **e**, Volcano plot depicting DEGs in EC2 endothelium in atheroma-containing segments compared to atheroma-free segments. **f**, Top 15 pathways that are more prevalent or less prevalent in EC2 in atheroma-containing compared to atheroma-free segments.
