## Supplementary Table 4 for "Coronary Artery Disease Transcriptomics Reveals Two Drivers of the Endothelial Cell SR-BI Expression and LDL Transport that Underlie Atherosclerosis"

| **Primer-probe for TaqMan Gene Expression Assay** | | | | |
| --- | --- | --- | --- | --- |
| **Gene** |  | **Cat. No** |  | **ID** |
| **mouse SR-BI** |  | **4351372** |  | **Mm00450226_m1** |
| **mouse HIF-1a** |  | **4331182** |  | **Mm00468869_m1** |
| **mouse ANGPTL4** |  | **4331182** |  | **Mm00480431_m1** |
| **mouse hprt** |  | **4331182** |  | **Mm00446968_m1** |
| **mouse VEGF** |  | **4331182** |  | **Mm00437306_m1** |
| **human SR-BI** |  | **4331182** |  | **Hs00969821_m1** |
| **human hprt** |  | **4331182** |  | **Hs01035168_m1** |
| **human Hif1a** |  | **4331182** |  | **Hs00153153_m1** |

**Supplementary Table 4- qPCR Primers, ChIP PCR Primers and CRISPR gRNA and Primers**

| **Primers for Sybr Green qPCR** | |  |
| --- | --- | --- |
| **Gene** | **Forward** | **Reverse** |
| **mouse β-actin** | **GATCAAGATCATTGCTCCTCCTG** | **AGGGTGTAAAACGCAGCTCA** |
| **mouse GAPDH** | **TGCACCACCAACTGCTTAGC** | **GGCATGGACTGTGGTCATGAG** |
| **mouse Ubiquitin** | **AGTGACGAGAGGCTTTGTCC** | **CGAAGATCTGCATTTTGACCTGT** |
| **mouse SR-BI** | **GCTGCGCTCGGCGTTGTCAT** | **GGGACGGGGATCTCCTTCCA** |
| **mouse KLF2** | **CCTTCGGTCTTTTCGAGGAC** | **TAAGGCTTCTCACCTGTGTGTG** |
| **human Hif1a** | **GGCCTCTGTGATGAGGCTTAC** | **CACCATCATCTGTGAGAACCA** |
| **human VEGF** | **ACAAATGTGAATGCAGACCAAA** | **CACCAACGTACACGCTCCA** |
| **human scarb1** | **ACTTCTGGCATTCCGATCAGT** | **ACGAAGCGATAGGTGGGGAT** |
| **human RPL19** | **AAAACAAGCGGATTCTCATGGA** | **TGCGTGCTTCCTTGGTCTTAG** |

| **Primers for ChipPCR** | |  |
| --- | --- | --- |
|  | **Forward** | **Reverse** |
| **human VEGF** | **CCTCAGTTCCCTGGCAACATCTG** | **GAAGAATTTGGCACCAAGTTTGT** |
| **human SCARB1** | **CAGTGGTGAGCTCAGGGAGGC** | **TCACGGGTGAGGCCGGATCTG** |

| **CRISPR gRNA Sequences** | |  |  |
| --- | --- | --- | --- |
|  | **Upstream** | **Downstream** | **catalog number** |
| **sgRNA1** | **TTCATGACGTGTTCTTGAAC** | **GCCCTTCCTGAATTTCTGTG** | **VB230802-1416jgv** |
| **sgRNA2** | **aaccaccgttgccctatttt** | **ATTTCTGTGTGGTCTAGGCT** | **VB230802-1425dkd** |

| **Primers for CRISPR Evaluation** | |  |
| --- | --- | --- |
|  | **Forward** | **Reverse** |
| **HIF1a binding site** | **CAGTGGTGAGCTCAGGGAGGC** | **TCACGGGTGAGGCCGGATCTG** |
| **Confirmation primers** | **GGCCAGGGTCAAATTCCAACATGC** | **AATGCAGTGGCACTGGCCCACAGC** |
| **Negative Control 1** | **CGGGGACATGGAAGTGTGCTCAGC** | **CGCAGCGTGCGTCCAAACGTCCTTC** |
| **Negative Control 2** | **TGGATTTGGTCCACAGGCTGTAGC** | **GGCTGGGCACTGCCTTGGTAAAGG** |
| **Negative Control 3** | **GATGAGTGGGACTTCCTCTCCACC** | **AGAAATGCTTTTGCAAGCCGGTGG** |
